# Gain of function mutagenesis through activation tagging identifies *XPB2* and *SEN1* helicase genes as potential targets for drought stress tolerance in rice

**DOI:** 10.1101/2020.05.13.092643

**Authors:** Mouboni Dutta, Mazahar Moin, Anusree Saha, Achala Bakshi, P.B. Kirti

## Abstract

We have earlier reported on the development of an activation tagged gain-of-function mutant population in an *indica* rice variety, BPT-5204 (Moin et al. 2016). Screening of these gain of function mutants for water-use efficiency (WUE) followed by physiological analyses revealed the activation of two helicases, ATP-dependent RNA (*SEN1*) and DNA (*XPB2*) encoding unwinding proteins in two different mutant lines. In the current study, we examined the roles of these genes in stable activation tagged mutants of rice for drought stress responses. Transcript profiling of *SEN1* and *XPB2* showed their significant up-regulation under various stresses (particularly ABA and PEG). The *SEN1* and *XPB2* tagged mutants exhibited reduced leaf wilting, improved revival efficiency, high chlorophyll and proline contents, profuse tillering, high quantum efficiency and yield-related traits in response to simulated drought (PEG) and hormone (ABA) treatments with respect to their controls. These observations were further validated under greenhouse conditions by periodic withdrawal of water. Germination of the seeds of these mutant lines indicates their ABA insensitivity under high ABA concentration. Also, the associated high up-regulation of stress-specific genes suggests that their drought tolerance might have been because of the coordinated expression of several stress responsive genes in these two mutants. Altogether, our results provided a firm basis for *SEN1* and *XPB2* as potential candidates for manipulation of drought tolerance and improving rice performance and yield under limited water conditions.

## 1. Introduction

Stresses like drought, salinity, extreme temperatures and fungal, viral or bacterial infections are serious threats to sustainable agricultural productivity. Hence, identification of genes responsible for orchestrating plant tolerance to the various stresses is a continuous process and a necessary step in developing tailored crop varieties, which can withstand such challenging and aggressive environmental conditions.

Helicases are molecular ATPases, which utilize the energy released during ATP hydrolysis to carry out their wide range of functions either on DNA (known as DNA helicases) or on RNA (known as RNA helicases). In addition to their housekeeping functions of being associated with inducing conformational changes in DNA or RNA, they are also reported to be involved in combating biotic and abiotic stress conditions in plants (Tuteja, 2003; Linder and Owttrim, 2009). A majority of the helicases have a three dimensional conserved core region consisting of two tandemly placed RecA domains (RecA1 and RecA2) connected via a flexible linker region (Sloan and Bohnsack, 2018). The core domain consists of about 350-400 amino acids and 14 conserved motifs serving as their catalytic pockets (Umate et al., 2010; Passricha et al., 2018). The functional diversity of helicases may arise either due to differential binding patterns of the nucleic acids or from the variations in their N or C terminal domains (Jankowsky and Fairman, 2007; Seraj et al., 2018). Based on the structural and functional characteristics, helicases are divided into six super families (SF1 to SF6). Of them, a majority of the DNA and RNA helicases fall under SF1 and SF2 category. SF1 family of helicases are well characterized and SF2 is considered to be the largest family of helicases (Seraj et al., 2018; Passricha et al., 2018).

Rice XPB2 (Xeroderma Pigmentosa group B2) is a DNA helicase (3′ to 5′ helicase) belonging to superfamily 2 group. The homologs of rice *XPB2* in yeast, Arabidopsis and humans are known as *RAD25 (SSL2), XPB2* and *XPB* (*ERCC3*), respectively (Bhatia et al., 1996; Umate et al., 2010). Any distortions in the DNA is repaired via nucleotide excision repair (NER) mechanism so that the damage is not passed on to the next generation (Guzder et al., 1995; Morgante et al., 2005). XPB2 is a subunit of eukaryotic transcription factor, TFIIH that opens up a DNA bubble during RNA polymerase II mediated transcription initiation (Bhatia et al., 1996, Morgante et al., 2005). It acts as a DNA-dependent helicase, which also helps in NER by unwinding the DNA at the site of lesion (Richards et al., 2008; Raikwar et al., 2015). Ergo, TFIIH in eukaryotes has a dual role *viz*. transcription initiation and DNA damage repair via nucleotide excision (Bhatia et al., 1996; Costa et al., 2001). Defects in XPB2 have been associated with an autosomal recessive disease Xeroderma Pigmentosum in humans. These mutants are more sensitive in response to photoperiod (Park et al., 1992; Costa et al., 2001). Recently, it has been reported that the promoter of rice *XPB2* gene is a multi-stress inducible one playing an important role in orchestrating plant stress tolerance (Raikwar et al., 2015).

Rice *SEN1* (t-RNA splicing endonuclease) is a RNA helicase belonging to the Upf1-like subfamily under superfamily 1B group, which unwinds the RNA in 5’ to 3’ direction. The homologs of rice SEN1 found in Arabidopsis, yeast and human are UPF1, SEN1 and SETX (*Senataxin*), respectively (Umate et al., 2010; Martin-Tumasz and Brow, 2015). The exact mechanism of rice SEN1 activity is yet to be understood clearly, but since the helicase domains are highly conserved among the eukaryotes (Han et al., 2017; Leonaite et al., 2017), these proteins might function in a manner similar to other reported eukaryotes.

In yeast, transcription termination of non-coding RNAs occurs via NNS (NRD1-NAB3-SEN1) complex (Sariki et al., 2016; Leonaite et al., 2017), where the NRD1-NAB3 heterodimer interacts with specific sequences on the nascent RNA (Han et al., 2017; Mischo et al., 2018) and with C terminal domain of the RNA polymerase II during termination (Mischo et al., 2011). This interaction helps in recruiting SEN1 onto the nascent RNA, which dislodges the RNA polymerase II by its helicase activity (Mischo et al., 2018). After transcription is terminated, the RNA is degraded by the combined activity of the TRAMP (TRF4/ AIR2/ MTR4 polyadenylation complex) complex and the exosomes (Leonaite et al., 2017; Mischo et al., 2018). SEN1 also plays an important role in several other functions like RNA processing, elimination of short protein coding sequences, resolving R loop structures and maintaining genomic stability (Mischo et al., 2011). Defects in SEN1 results in defective R loop resolution and an increase in its frequency (Martin-Tumasz and Brow, 2015; Leonaite et al., 2017; Mischo et al., 2018), genomic instability and defects in replication (Mischo et al., 2018). Apart from regulating the expression of non-coding genes, SEN1 also co-ordinates the expression of small protein coding genes like *NRD1, HRP1, IMD2* and *CYC1* (Steinmetz et al., 2006). Yeast cells with N-terminal truncation of SEN1 had higher cell death and shortened life span. Arabidopsis homologue of *SEN1, UPF1* plays an important role in nonsense mediated decay (NMD) of abnormal RNA. It helps the plant in maintaining proper seed size (Yoine et al., 2006), floral and vegetative development (Arciga-Reyes et al., 2006). Thus, it has significant roles in regulating both transcription and translation in most of the eukaryotes. SEN1 was also found to be involved in transcription coupled repair mechanisms. (Li et al., 2016). Hence, the basic function of SEN1 is likely to dissociate any stalled elongation complex on the nucleic acid during transcription.

In an earlier investigation, using tetrameric enhancer-based activation tagging system, we have generated a large population of gain-of-function mutant lines in a widely cultivated *indica* rice variety BPT-5204 (Samba Mahsuri) (Moin et al., 2016). Upon screening of these mutants under limited water conditions along with associated phenotypic and physiological studies, some of them showed high water use efficient (WUE) phenotypes indicating that the genes, which became activated in them through the integrated enhancers might have roles in improving the WUE in rice. We initially identified five mutant lines with high quantum efficiency and low Δ^13^C, which are the proxies for WUE, showing the activation of transcription factors (GRAS and WRKY 96) and proteins involved in protein ubiquitination (cullin4) and ribosome biogenesis (RPL6 and RPL23A) (Moin et al., 2016). Subsequent analysis of sequences flanking the activation tags (4X enhancers) in two other WUE mutants resulted in the identification of two plant RNA and DNA helicases, *SEN1* and *XPB2*, respectively. In the present study, we characterize the roles of these two genes in response to various stress conditions, particularly drought with an emphasis on seed yield and productivity apart from WUE in *indica* rice. Our findings suggested that these gene encoded proteins, in addition to their basic cellular housekeeping activities (nucleic acid unwinding) also have roles in stress responses by possibly maintaining the genomic integrity of the plant upon the onset of environmental stresses.

## 2. Materials and Methods

### 2.1 Identification of *SEN1* and *XPB2* helicases in activation tagged mutants

The *Ac/Ds*-based activation tagging vector, pSQ5 (Qu et al., 2008) was used to generate gain-of-function mutant lines of *indica* rice cultivar, Samba Mahsuri (variety BPT-5204). The *Ds* element of the vector consists of tetrameric repeats of CaMV35S enhancers (4X enhancers), which when integrated in the plant genome, upregulate the genes present 10Kb upstream or downstream from the point of their integration in the genome. This vector was transformed into rice using *in planta* and callus-mediated rice transformation protocols and the transformed plants were confirmed by molecular analysis (PCR and Southern-blot hybridization) and screening on solid MS medium containing 50 mg/L Hygromycin (Moin et al. 2016). Because the T-DNA of transformed vector contains both the *Ac* and *Ds* elements, four types of progeny plants are expected with respect to their genome composition as a result of meiotic segregation. These include the mutants containing both the elements (*Ac*^+^/*Ds*^+^) or *Ac* alone (*Ac*^+^/*Ds*^-^), *Ds* alone (*Ac*^-^/*Ds*^+^) and null (*Ac*^-^/*Ds*^-^). Among these, the *Ac*^-^/*Ds*^+^ mutants does not undergo transposition because of the absence of *Ac* and also exhibit stable expression of the enhancer-tagged genes and therefore, we selected only such mutants. The strategy for selection of stable *Ds* mutants from the segregating population was followed using antibiotic (*hpt*II) screening as described earlier (Moin et al. 2016).

Once the transgenic nature of the mutants was confirmed and stable lines were selected, the transformed plants were carried forward to further generations and the identified mutant lines were investigated for WUE trait by growing them under limited water conditions followed by phenotypic and physiological studies related to WUE. Carbon isotope discrimination (Δ^13^C) is a non-invasive way of determining the WUE in plants. The negative relationship between WUE of a plant and Δ^13^C values is an effective way of identifying plants with an improved efficiency under limited water conditions. (Chen et al., 2011) In C_3_ plants, the molar abundance ratio (R) of ^13^C/^12^C is less than that of atmosphere because of the discrimination of the plant in uptake of ^13^C during photosynthesis. The carbon isotope composition (d^13^C‰) of plants is calculated by the formula [(R_sample_/R_standard_)-1]×10^3^ and is compared with the standard of Pee Dee Belemnite (PDB) fossil carbonate. The carbon isotope discrimination (Δ^13^C‰) is measured by the formula [(d^13^C_a_-d^13^C_p_)/(1+ d^13^C_p_)×10^3^ where d^13^C_a_ and d^13^C_p_ are the d^13^C values of atmosphere and plant respectively. (Farquhar et al., 1982, Farquhar et al., 1989,Gao et al., 2018). For calculating the carbon isotope discrimination values, 500 mg of mature leaf samples of WT, XM3 and SM4 lines grown under limited water conditions were collected and dried at 65°C for 3 days. The samples were powdered and the carbon isotope was measured using Isotope Ratio Mass Spectrometer (IRMS).

Based on their performance under limited water conditions, a few mutants that appeared to have high WUE were identified. Further, to identify the corresponding gene responsible for this trait, a flanking sequence analysis using TAIL-PCR (thermal asymmetric interlaced PCR) was performed on these mutants that exhibited enhanced WUE using one degenerate primer and three nested primers, which were 1 Kb, 500 bp and 100 bp upstream to the 5’ end of the *Ds* element. The protocols for the TAIL-PCR and the related analyses were essentially followed as per Moin et al. (2016). Three rounds of PCR was conducted in which the product of each round was diluted 50-100 fold depending on the band intensity observed on 1% agarose gel and 1 µl of the diluted sample was used as template for the next round of amplification. The protocol followed was the same as used in Moin et al. (2016). The final amplicons were cloned in pTZ57R/T and sequenced. The identification of site of enhancer integration in the rice genome was done by subjecting the sequence obtained to BLAST search in the rice genome database (RGAP-DB). Two of these potential mutants with high WUE were found to have *SEN1* and *XPB2* helicase genes within the 20 kb region of enhancer integration. The activation tagging of these helicase genes was analyzed using semi-quantitative and qRT-PCR using appropriate primers.

### 2.2 Quantitative PCR (qRT-PCR) analysis of flanking genes

The two mutant lines *viz*. En.124 (named as SM4) and DEB.36 (named as XM3), which exhibited normal growth and yield-related parameters under limited water conditions with respect to WT were selected for semi-quantitative PCR analysis. The transcript levels of all the genes present in a region of 10 kb upstream and downstream (total 20 kb region) from the site of enhancer integration in the rice genome were studied. A total of 5 genes in En.124 and 3 genes in DEB.36 were situated in a total of 20 kb enhancer integration region. The levels of transcripts of the tagged genes were compared with the wild type (WT) plants and was further validated by quantitative real time PCR (qRT-PCR) using gene specific primers in three biological and technical replicates. The total RNA was isolated from the leaves of 60 days-old plants using Tri-Reagent (Takara Bio, UK) and 2 µg cDNA (Takara, Clonetech, USA) was prepared as per the manufacturer’s protocol. The cDNA was diluted seven times and 2 µl of it was used for qRT-PCR. The expression level was normalized using rice *actin* (*act1*) as an internal reference gene and the fold change was calculated using ΔΔC_T_ method (Livak and Schmittgen, 2001).

### 2.3 *In silico* promoter analysis of *SEN1*

About 1 Kb upstream sequence from start codon of *SEN1* (LOC_Os10g02930) was retrieved from RGAP-DB and was subjected to an *in-silico* analysis for the presence of *cis*-acting elements using the PlantCARE (Lescot et al. 2002) online tool. Similar promoter analysis of XPB2 gene (LOC_Os01g49680) has been reported earlier (Raikwar et al., 2015).

### 2.4 Differential transcript analysis of *SEN1* and *XPB2* helicases

To understand the involvement of *SEN1* and *XPB2* genes in response to biotic and abiotic stresses, their differential transcription patterns were analyzed in 10 day old seedlings of WT BPT-5204 rice. For this, various plant hormones and abiotic stress-inducing agents like 2 mM salicylic acid (SA), 100 μM methyl jasmonate (MJ), 15% polyethylene glycol (PEG 8000), 250 mM sodium chloride (NaCl), 100 μM abscisic acid (ABA) and heat at 42 °C were applied to the seedlings. Salt, simulated drought and phytohormone treatments were given by dipping the seedlings in their respective solutions, while heat treatment was induced by exposing the seedlings in a hot air oven maintained at 42 °C. The root and shoot samples were collected immediately after the onset of the stress treatments denoted separately as the 0 hr, followed by 3 hr, 6 hr, 12 hr, 24 hr and 48 hr after treatments. Seedlings grown in stress-free medium under similar growth conditions were used as control for normalization.

For the transcript analysis of these genes under biotic stress conditions, leaf samples of rice infected with *Xanthomonas oryzae* pv. *oryzae* (*Xoo*, causing bacterial leaf blight) and *Rhizoctonia solani* (causing sheath blight) were used. The infection process of these pathogens to rice were followed as described earlier (Moin et al. 2016b, Saha et al. 2017). Leaf samples from untreated plants were used as control to normalize the expression. Rice *actin* was used as the internal reference gene. All qRT-PCR experiments were performed by using gene specific primers in three technical and three biological replicates. The protocol for cDNA preparation and fold change calculation is same as mentioned earlier.

### 2.5 Measurement of growth and phenotypic parameters of the activation tagged mutant lines

The two stable mutants with activated *XPB2* and *SEN1* helicase genes were selected to study their behavior in the presence of abiotic stresses. For this, seeds of XM3, SM4 and WT (BPT 5204) were surface sterilized with 70% ethanol for 50 to 60 sec followed by 4% sodium hypochlorite for 15 to 20 min and final wash with sterilized double-distilled water (Saha et al. 2017). They were then germinated on Murashige and Skoog medium for 25 days. Subsequently, they were transferred to test tubes containing half strength of liquid nutrient Yoshida solution (Yoshida et al., 1976) along with stress-inducing agents. Simulated drought stress (10% of PEG-8000) and phytohormone treatment (50 and 75 µM of ABA) were applied to the seedlings. Seedlings kept in plain half-strength liquid Yoshida solution were served as control.

The seedlings were subjected to fresh stress by changing the solutions after every 7 days. The seedlings were transferred to full strength Yoshida liquid medium twenty Days After Stress (DAS) treatment for revival experiments. Before shifting, root and shoot samples were collected separately for transcript analysis and the whole seedling samples were collected for biochemical experiments. The samples were freeze-dried in liquid nitrogen and stored at -80°C. Similar pattern was followed during revival of the plants. The samples were collected for biochemical experiments twenty Days After Revival (DAR) and the seedlings were transferred to the greenhouse for checking their yield parameters and photosynthetic efficiency. The plants were acclimatized using similar greenhouse conditions as discussed earlier. The root length (cm), shoot length (cm), fresh weight (g), percentage of leaf wilting and survival of the seedlings were constantly recorded. In order to understand the leaf wilting percentage, we measured the length of whole leaf and the wilting region separately and calculated the percentage of wilting of each leaf. The mean values are provided as a line graph.

The revived seedlings were transferred to greenhouse and were allowed to acclimatize for thirty days. During their growth, various parameters were measured which included; plant height, number of total and productive tillers (panicles), boot leaf length v/s panicle length, and yield related parameters like number of branches per panicle, number of seeds per panicle and per plant and weight of 100 seeds.

### 2.6 Measurement of chlorophyll fluorescence and Carbon isotope composition

The chlorophyll present in the reaction centre of photosystem II (PSII) is responsible for absorbance of light energy. A part of the absorbed energy is re-emitted as fluorescence, which is the same as the one utilized for photosynthesis. Hence, a measurement of chlorophyll fluorescence connotes the quantum yield of a plant. Photosynthetic performance of plants is measured indirectly by a Pulse Amplitude Modulated fluorometer (PAM). It measures the activity of the PSII by exposing a dark adapted leaf to a high pulse of light. The minimal level of fluorescence observed upon irradiation is considered as F_o_, while F_m_ indicates the maximum value of fluorescence obtained after application of saturated pulse of light. The difference between F_o_ and F_m_ is F_v_ or the variable fluorescence. The PSII activity is measured by the ratio of the variable fluorescence and the maximum fluorescence (Fv/Fm). Thus, F_v_/F_m_ ratio is indicative of overall photosynthetic efficiency and also the level of stress experienced by a plant under unfavourable conditions. In a healthy unstressed plant, the F_v_/F_m_ value ranges around ∼0.83 (Murchie and Lawson, 2013). This value corresponds to maximum photosynthetic yield in a plant. A significant reduction in this value (less than 0.83) indicates induction of stress in the plant. In our experiments, we have used a portable MINI-PAM and followed the manufacturer’s protocol (Walz, Effeltrich, Germany) to study the quantum efficiency of selected mutants with respect to their corresponding controls. All the plants were initially incubated in dark for 30 minutes and were then subjected to a short pulse of 8000 μmol m^-2^ s^-1^ light. The F_v_/F_m_ ratio was plotted as a histogram.

### 2.7 Estimation of Chlorophyll and proline contents

Two sets of samples collected after application of stress followed by their revival were used for chlorophyll estimation. About 100 mg of tissue was ground to a fine powder in liquid nitrogen and chlorophyll was extracted using di-methyl sulphoxide (DMSO). The corresponding absorbance was taken at 663 nm and 645 nm using an UV spectrophotometer and chlorophyll a, b and total chlorophyll contents were calculated (Zhang et al, 2009). The mean and standard error was plotted in the form of a bar graph.

Similar sets of samples were used for estimation of contents of important osmolyte, proline (Bates et al. 1973). For this, 100 mg of plant tissue was homogenized in 5 ml of 3% sulphosalicylic acid. The homogenate was centrifuged at 12,000 rpm for 15 min and the supernatant was used for proline estimation. The supernatant was mixed with acid ninhydrin and glacial acetic acid (400 µl each) and was incubated at 100 °C for one hour. The solution was immediately transferred to an ice bath, cooled, and 800 µl toluene was added and the mixture was vortexed. Subsequently, the organic phase was pipetted out to a new tube and the absorbance was recorded at 520 nm. The proline content was estimated by using a standard curve and the values from individual samples were indicated as a bar diagram.

### 2.8 Pot-level water withholding treatments

To study the response of *SEN1* and *XPB2* expressing activation tagged mutants under pot-level drought conditions, fifteen day-old mutant and WT seedlings grown under culture room conditions were shifted to the pots containing black alluvial soil provided with ample water and maintained under greenhouse conditions (30 ± 2 °C, 16 h light/ 8 h dark photoperiod). After thirty days, the overlay water from the pot was removed and all the WT and mutant plants were subjected to drought conditions in separate pots for three and seven consecutive days. In our previous study, we noticed that the Permanent Wilting Point (PWP) for BPT-5204 in our greenhouse conditions was 21 days (Moin et al. 2017). The drought experiments are usually defined in Percent Field Capacity (FC), which mainly depends on the type of soil. For black alluvial soils available in South India, which was also used in the current study, the PWP occurs between a FC of 10 to 18% (http://www.indiawaterportal.org). Accordingly, in the present study, the FC on three and seven day drought treatments might be ∼60 and 40%, respectively. After completion of drought treatments, plants were uprooted and their root and shoot samples were collected separately for studying the transcript patterns of potential drought responsive genes under stress conditions. A few plants were then allowed to revive after the drought treatments by gradually supplying sufficient amount of water as required for normal rice cultivation and were grown till plant maturity. Comparative yield-related studies were made between treated mutant plants with corresponding treated and also untreated control counterparts to study whether the total yield of mutants was decreased, sustained or improved under drought.

### 2.9 Transcript analysis of stress-responsive genes

The root and shoot tissues of mutant and WT plants, which were exposed to phytohormone, (ABA), drought-inducing agent-PEG and pot-level drought (both three and seven day old samples) were used to check the transcript pattern of seven stress regulated genes such as *OsTPP1, OsLEA3-1, OsPP2C, OsDREB2B, OsNAC1, OsNAC2* and *OsSIK1*. These genes are known to be involved in ABA signaling and in modulating heat, cold, salt and drought stresses. The transcript levels of these genes in tagged lines were normalized with those of the corresponding stress-induced WT plants. About 2 µg concentration of cDNA was prepared, diluted 10 times and 2 µl of it was used for qRT PCR..

### 2.10 Seed germination assay

The seeds of the tagged lines along with WT were surface sterilized and allowed to germinate on half strength MS media containing 50 µM and 75 µM ABA under 16/8 hr light/dark cycles. Seeds germinated on half strength MS media without ABA were used as untreated controls. The germination of the seeds was documented after 5 days.

### 2.11 Statistical analysis

All the qRT-PCR experiments were conducted in three biological and technical replicates, whereas phenotypic and physiological experiments were conducted in replicates of three plants. Statistical analysis of the mean values have been calculated using one way ANOVA in SigmaPlot v11 at a significance level *P*<0.001 marked as “a”, *P*<0.025 marked as “b” and *P*<0.05 marked as “c” in bar diagrams.

## 3. Results

### 3.1 Identification of tagged genes

Out of the selected activation tagged mutant lines identified for enhanced WUE through studies on photosynthetic performance and Δ^13^C analysis that could be correlated with high WUE, two lines SM4 and XM3 were chosen for a detailed analysis. This is because although the roles of these two gene encoding proteins have been well studied in nuclear activities but not been emphasized in stress responses or WUE. It was observed that under limited water conditions, WT had higher Δ^13^C (23.75‰) values than both XM3 (20.05‰) and SM4 (20.14‰) mutant lines, respectively (Fig. S1). Since lower carbon discrimination values is an indication of higher WUE (Δ^13^C is inversely related to WUE), these lines were carried further for flanking gene analysis. TAIL-PCR analysis was performed on the selected mutants, SM4 (En.124) and XM3 (DEB.36) to confirm the transgenic nature of the selected plants, fix the site of integration of the tetrameric enhancers in their genomes and identify the genes in their vicinity. The list of TAIL-PCR primers are provided in Stable 1. The sequences obtained from TAIL-PCR were subjected to a nucleotide BLAST in the rice genome database to map the exact location of the point of integration of the tetrameric enhancer element (Fig. 1a & b). This led us to identify the genes located on either sides of the 4X enhancers in two mutants.

**Table 1:** Comparative analysis of yield related traits at pot-level drought conditions. Chart showing phenotypic characteristics observed in the tagged lines and the WT plants post imposal 3 days and 7 days of drought and revived till seed setting. The observations included number of primary branch/panicle, numbers of seeds/branch of the panicle, seeds/panicle, total seeds/plant. The mean ± standard error is represented in the chart.

| Treatment | Parameter | WT | XM3 | SM4 |
| --- | --- | --- | --- | --- |
| UT | Primary branch/panicle | $7.62 \pm 0.37$ | $6.88 \pm 0.38$ | $6.87 \pm 0.29$ |
| | Seeds/branch | $16.86 \pm 0.99$ | $16.29 \pm 2.03$ | $17.35 \pm 1.23$ |
| | Seeds/panicle | $112.4 \pm 7.08$ | $102 \pm 3.08$ | $104 \pm 5.72$ |
| | Total seeds/plant | $322 \pm 23$ | $299.5 \pm 8.5$ | $324.5 \pm 24.5$ |
| 3D drought | Primary branch/panicle | $6.11 \pm 0.42$ | $8.22 \pm 0.27$ | $7.33 \pm 0.64$ |
| | Seeds/branch | $14.92 \pm 1.23$ | $16.08 \pm 0.96$ | $15.55 \pm 1.02$ |
| | Seeds/panicle | $108.16 \pm 8.29$ | $124.14 \pm 2.98$ | $111 \pm 5.19$ |
| | Total seeds/plant | $306.66 \pm 14.83$ | $384.5 \pm 11.5$ | $338.5 \pm 22.5$ |
| 7D drought | Primary branch/panicle | $6.66 \pm 0.49$ | $7.42 \pm 0.29$ | $7 \pm 0.32$ |
| | Seeds/branch | $13.83 \pm 0.85$ | $14.27 \pm 1.16$ | $16.41 \pm 0.84$ |
| | Seeds/panicle | $87.83 \pm 11.30$ | $104.75 \pm 6.73$ | $106.14 \pm 4.17$ |
| | Total seeds/plant | $200 \pm 27.59$ | $414 \pm 83.76$ | $377.33 \pm 8.45$ |

**Figure 1:**
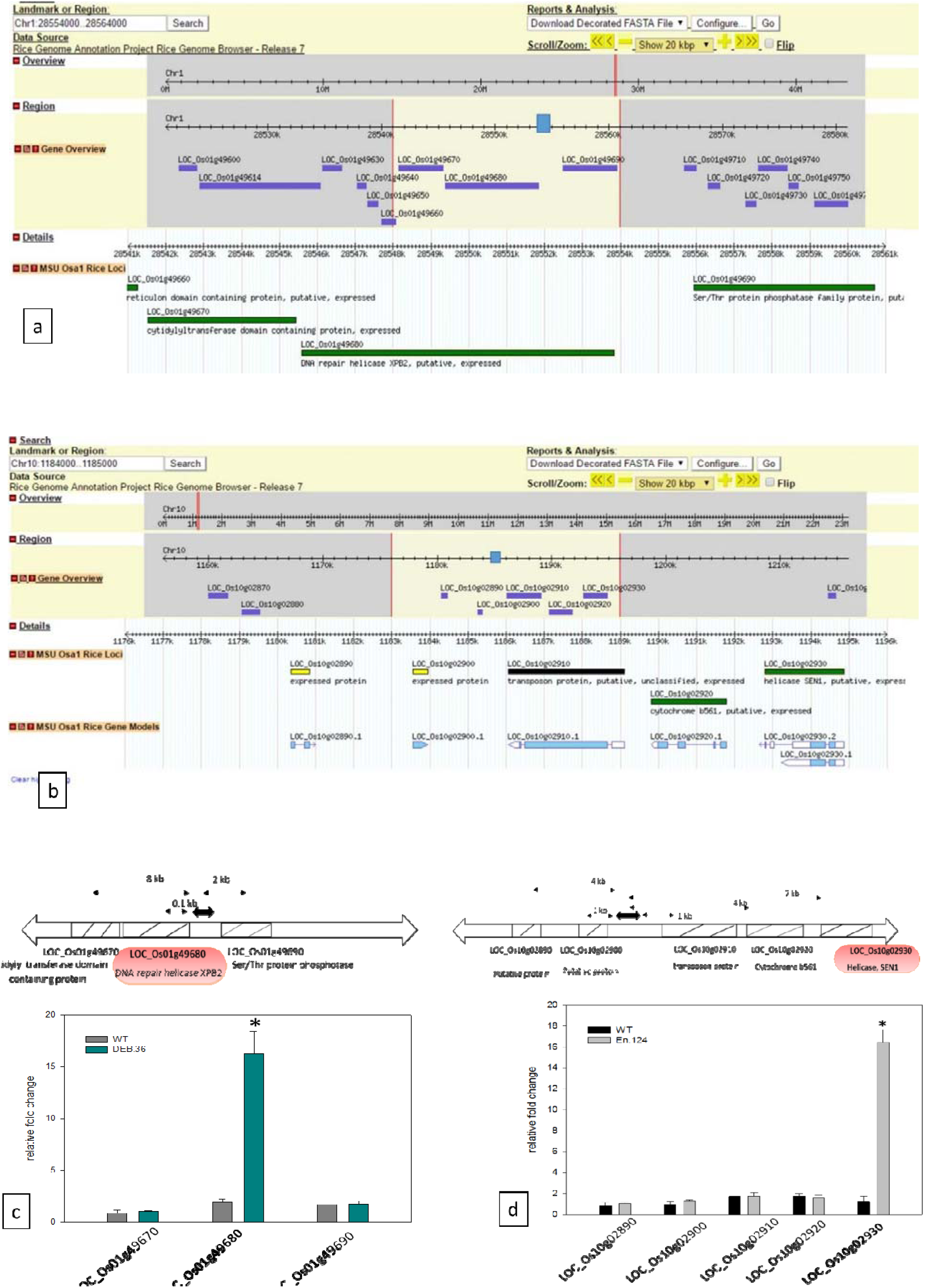
Gene map and quantitative real-time PCR of the tagged genes. Gene map locating the point of integration of the tetrameric enhancer element of activation-tagging vector and its 10 kb upstream and downstream genes as obtained from rice genome database and the subsequent Quantitative real-time PCR (qRT-PCR) analyses of the tagged genes. The sequence obtained from TAIL-PCR of (a) XM3 (DEB.36) and (b) SM4 (En.124) was subjected to BLASTN analysis in the rice genome database. The blue rectangular box indicate the point of integration of the enhancer element and the selected region indicate the surrounding 20Kb region of the enhancers. In the line XM3, 3 genes were present (LOC_Os01g49670, LOC_Os01g49680 and LOC_Os01g49690) and in the line SM4 (En.124), 5 genes were present (LOC_Os10g02890, LOC_Os10g02900, LOC_Os10g02910, LOC_Os10g02920, LOC_Os10g02930) in the selected region. (c), (d): Quantitative real-time PCR (qRT-PCR) analyses showed up to 16 fold upregulation of two genes i.e. XPB2 and SEN1 as compared to the wild type plant in XM3 and SM4 lines respectively. Other tagged genes showed an expression level similar to that of the wild type. The data was normalized using rice actin as an internal reference gene. One way ANOVA was performed at a significance level P<0.05 annotated by *.

In the tagged line, XM3, LOC_Os01g49670, LOC_Os01g49680 and LOC_Os01g49690 were present in a 20 Kb region. Among these, LOC_Os01g49670 and LOC_Os01g49680 were situated 8 Kb and 0.1 Kb upstream of enhancers, encoding cytidylyl transferase domain containing protein and the DNA repair helicase XPB2, respectively (Fig. 1b). The LOC_Os01g49690 was located 2 Kb downstream from the enhancer integration and encodes a serine/threonine protein phosphatase. In SM4 mutant, the enhancers were flanked by LOC_Os10g02890, LOC_Os10g02900, LOC_Os10g02910, LOC_Os10g02920 and LOC_Os10g02930 loci in a 20 kb region. The first two genes that were located 4 Kb and 1 Kb upstream of enhancers encodes unidentified putative proteins. The remaining three that were located 1 Kb, 4 Kb and 7 kb downstream encodes a transposon protein, cytochrome B561 and SEN1 helicase, respectively.

A semi-quantitative PCR analysis (Fig. S2a & b) showed an upregulation of the genes, *XPB2* (LOC_Os01g49680) and *SEN1* (LOC_Os10g02930) in the lines XM3 and SM4, respectively. This was further validated via qRT-PCR analysis (Fig. 1c & d), which showed 16 fold upregulation of both the genes in their corresponding tagged mutants with respect to WT, while there was no considerable upregulation of other genes situated in the 20 kb regions.

### 3.2. Promoter analysis of *SEN1*

The *XPB2* promoter region was earlier reported to contain many *cis*-acting elements including the one for early responsiveness towards dehydration (ABRELATERD1), dehydration responsive element (CBFHV) and MYBCORE element, that is responsive to water stress (Raikwar et al., 2015). Our *in-silico* promoter analysis of *SEN1* revealed several *cis*-acting stress responsive elements. The exact location and the detailed list of the *cis*-regulatory elements is provided in the Supplementary Figure S2a and Table S2. This included 12 MYB binding elements (CAACTG and CAACCA/CAACAG) and two ABRE motifs (CGTGG) responsible for drought responsiveness, among others. The high transcriptional upregulation of these two genes in response to ABA and simulated drought inducing agent, PEG can be correlated with the presence of these corresponding responsive elements in their putative promoter regions.

### 3.3. Transcript analysis of *XPB2* and *SEN1* under biotic and abiotic stresses

The *SEN1* and *XPB2* genes were found to be involved in improving the WUE of rice under limited water conditions, hence, we ascertained the responsiveness of these two genes to other stresses also. Their transcript patterns were studied in response to various abiotic stress inducing factors such as SA, MJ, NaCl, ABA, PEG and heat. It was observed that both the genes showed a significant upregulation, particularly in the root tissues as compared to the shoots. We have categorised the expression pattern of the two genes as early (expressing within 3 to 12h of treatment) and late (expressing after 12 h of treatment) responses.

In shoot, the level of *XPB2* transcripts was induced by more than 3 fold by SA, NaCl, ABA, PEG and high temperature (42°C) (Fig. 2a) as an early stress response which reached their peaks during this time period except PEG which increased up to 6 fold after 48h. The transcript level in roots was upregulated by more than 5 fold under all six imposed conditions within 3 to 12 h indicating an early responsiveness of the gene. In response to PEG, the maximum transcript level (10 fold) was maintained after 12h, while that under NaCl (11 fold), ABA (48 fold) and high temperature (14 fold) was observed after 48h (Fig. 2b). In shoots, *SEN1* transcript level was induced as an early response under ABA (9 fold) and PEG (3 fold) treatment within 3h. Later, it decreased under ABA, while that under PEG reached a peak of 18 fold after 48h (Fig. 2c) An early pattern of response of *SEN1* was observed under SA, NaCl, ABA and PEG in roots, while a late response was observed under MJ and high temperature. The transcript level was maintained under PEG (5 fold) from 12 to 48h, while that of ABA kept on increasing and reached its maximum up to 320 fold at the end of 48h (Fig. 2d).

**Figure 2:**
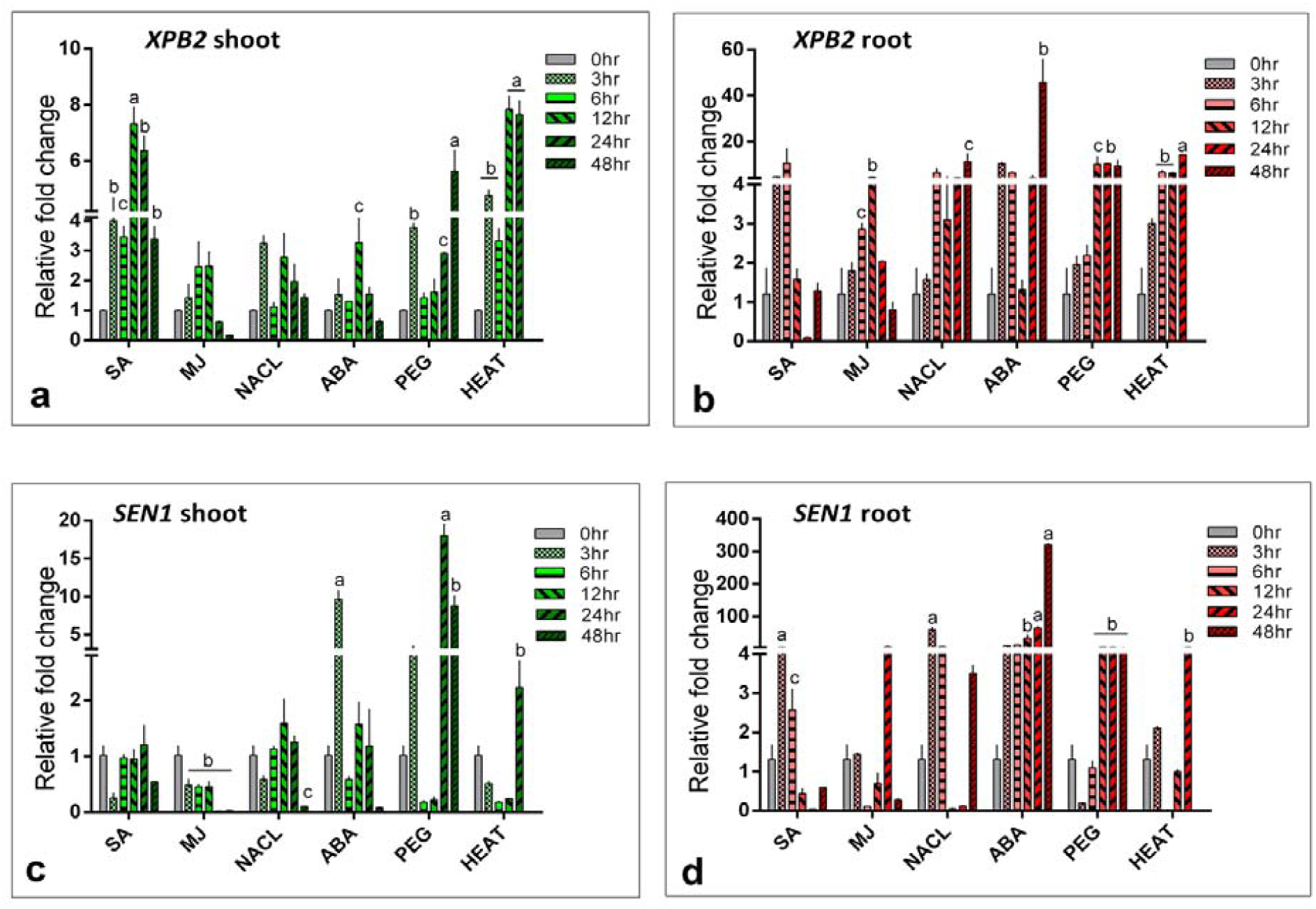
Transcript analysis of *XPB2* and *SEN1*. Quantitative real-time PCR (qRT-PCR) analyses of (a), (b) XPB2 shoot and root (c), (d) SEN1 shoot and root respectively in response to hormonal and chemical treatments. 10days old rice seedling were subjected to SA, MJ, NaCl, ABA, PEG and heat treatments and root and shoot tissues were collected at various time points. Rice actin was used as the internal reference gene. The fold change was calculated using ΔΔC_T_ method. The mean and the standard error is plotted in a vertical bar graph. One way ANOVA was performed at a significance level P<0.001 marked as a, P<0.025 marked as b and P<0.05 marked as c.

On the contrary, *Xoo* and *R. solani* pathogens failed to significantly induce the transcript levels of *SEN1* and *XPB2*. Upon infection with *Xoo*, both *SEN1* and *XPB2* were downregulated by 0.4 fold and 0.3 fold, respectively. In response to *R. solani*, the transcript level of *SEN1* was similar to that of untreated samples whereas, *XPB2* showed an elevation of around 2.4 fold (Fig. S4).

### 3.4. Phenotypic and physiological analyses of the tagged mutants under PEG and ABA

Post twenty DAS, the two tagged mutants exhibited better tolerance towards simulated drought (10% PEG) and phytohormone (50 µM and 75 µM ABA) treatments. The wilting of WT ranged between 9 (50µM ABA) to 60% (PEG) cumulatively under all three conditions and 30 (PEG) to 60% (50µM ABA) recovery post twenty DAR. (Fig. S4a to d). XM3 and SM4 lines, on the contrary, exhibited maximum wilting of 12 and 14% under PEG and a maximum recovery of 85 to 95% under ABA 50µM and PEG post twenty DAR, indicating that the enhanced expression of both the helicases rendered the plants to be more drought tolerant than the WT.

#### 3.4.1 Seedling shoot and root parameters

The fresh weight, shoot and root lengths (Fig. S5a to c) of the tagged lines were recorded 10 and 20 DAS. Post twenty DAS, XM3 lines had higher fresh weight (0.16 to 0.21 g) because of increased branching but showed similar shoot and root lengths compared with the corresponding WT (0.11 to 0.17 g). In SM4 lines, the fresh weight was observed to be 0.21 to 0.25 g due to longer shoot and root lengths than the WT. The mean observations of fresh weight, shoot and root lengths of the WT and mutant plants 10 and 20 DAS were plotted as a bar graph.

#### 3.4.2 Yield-related traits

Post acclimatization in the greenhouse, the XM3 and SM4 lines were observed to have better phenotypic parameters with increased number of tillers and panicles per plant, improved plant height, boot leaf, panicle length, number of seeds per plant, seed weight and photosynthetic efficiency (Fig. 3a & b)). The histogram depiction of the phenotypic parameters are provided in Figure S6 (a to c) and the mean values along with their standard errors are provided in Supplementary Table S3.

**Figure 3:**
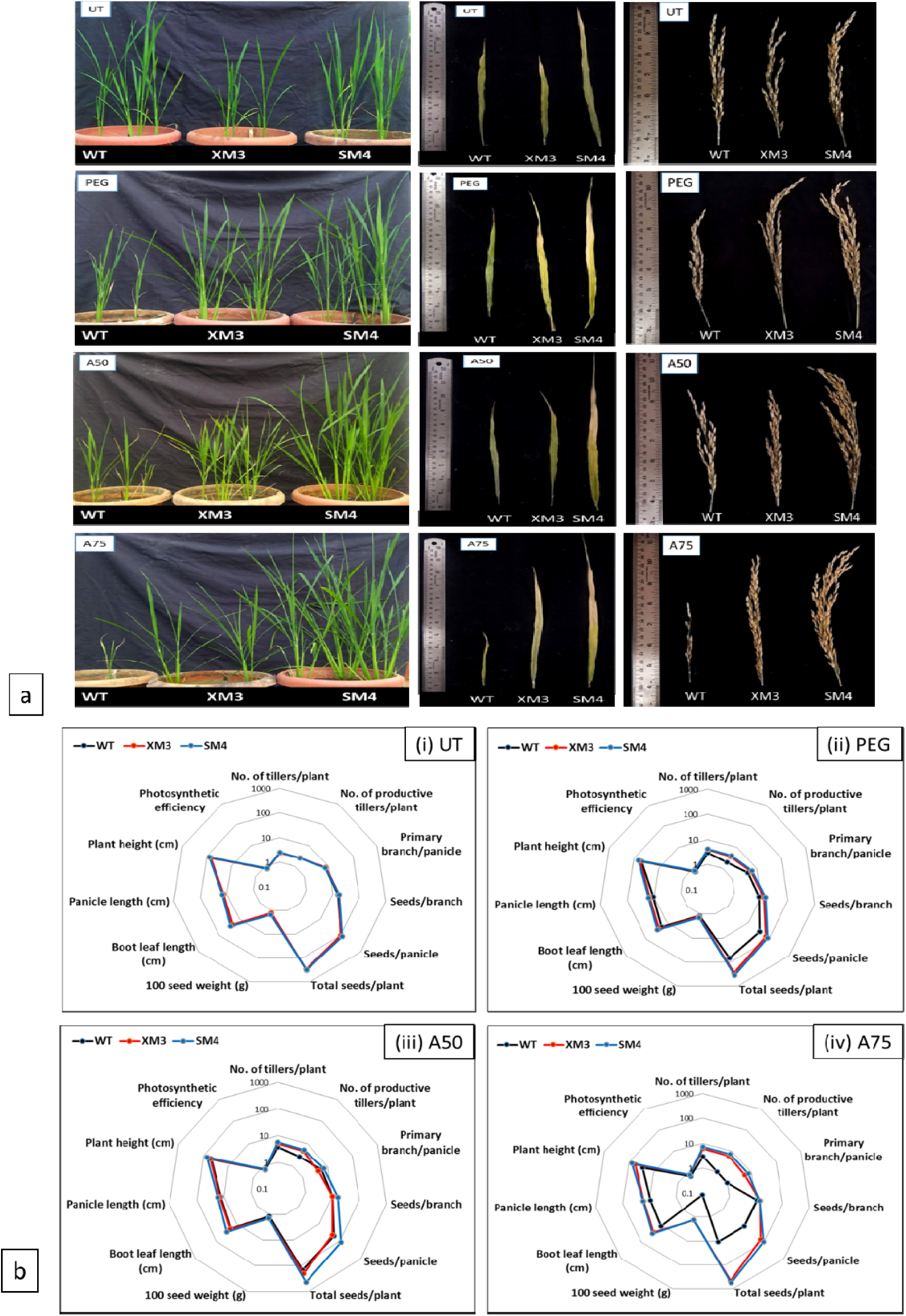
Physiological analysis of tagged lines. Phenotypic and physiological observations done post acclimatization of the tagged lines (XM3 and SM4) in comparison with the WT plants. These include the difference in the plants height, boot leaf and panicle length (a) as observed phenotypically. (b) Represents the observed phenotypic features plotted as a radar graph. The parameters plotted are plant height (cm), number of tillers/plant, number of productive tillers/plant, primary branch/panicle, seeds/branch, seeds/panicle, total seeds/plant, 100 seed weight (g), boot leaf length (cm) and the panicle length (cm) and photosynthetic efficiency of the WT lines in comparison to XM3 and SM4 lines. (i), (ii), (iii) and (iv) represent different conditions such as untreated, 10% PEG, 50µM and 75µM ABA respectively. The mean values have been plotted in a logarithmic (log_10_) scale.The tagged lines were observed to perform better under simulated stress conditions than the WT lines. The decrease in size of the black undecagon (WT) represents the same.

The WT revived from all three stress treatments had 2 to 3 tillers per plant with 1 to 2 bearing panicles, shorter plant height, boot leaf and decreased panicle length. XM3 and SM4 had 3 to 7 tillers and 3 to 9 tillers per plant, respectively with all being productive. These were taller, had longer boot leaf and panicle lengths than the WT. The boot leaf and panicle lengths were observed to be codependent.

In response to 75 µM ABA, a single WT plant survived bearing only 11 seeds. The total seed yield of XM3 (∼461 seeds) and SM4 (∼557 seeds) were 97 and 98% more than corresponding WT, respectively under 75 µM ABA treatment. A 75 to 80% (XM3 and SM4 respectively) difference in seed yield was also observed under 10% PEG. Under untreated conditions, the weight of 100 seeds were almost similar in WT and the tagged lines, however, a significant difference was noted under PEG and ABA 50µM stress conditions (both PEG and 50µM ABA. These physiological characters indicated a decreased sensitivity of SEN1 and XPB2 towards high ABA concentrations.

#### 3.4.3. Photosynthetic efficiency

In untreated WT, the quantum yield was found to be 0.74, whereas in both the untreated tagged mutants the efficiency was 0.77. Under 10% PEG, 50 and 75 µM ABA, the quantum yield of the WT decreased to 0.63 and 0.67, respectively. But the tagged lines continued to maintain high quantum efficiency which was almost similar to the untreated controls. The Fv/Fm ratio ranged from 0.72 to 0.77 in the tagged lines even after PEG and ABA treatments.

### 3.5. Chlorophyll and proline estimation

The contents of a, b and total chlorophyll were measured in the WT and the tagged lines post stress (Fig. 4a to c) and post recovery (Fig. S7a to c). In XM3 the chlorophyll a, b and total chlorophyll contents was 17 µg, 10 µg and 27 µg/50 mg fresh weight, respectively compared with WT which had 10 µg, 8 µg and 21 µg/50 mg fresh weight, respectively. The chlorophyll content of XM3 post recovery was observed to be moderately high under PEG and 50µM ABA.

**Figure 4:**
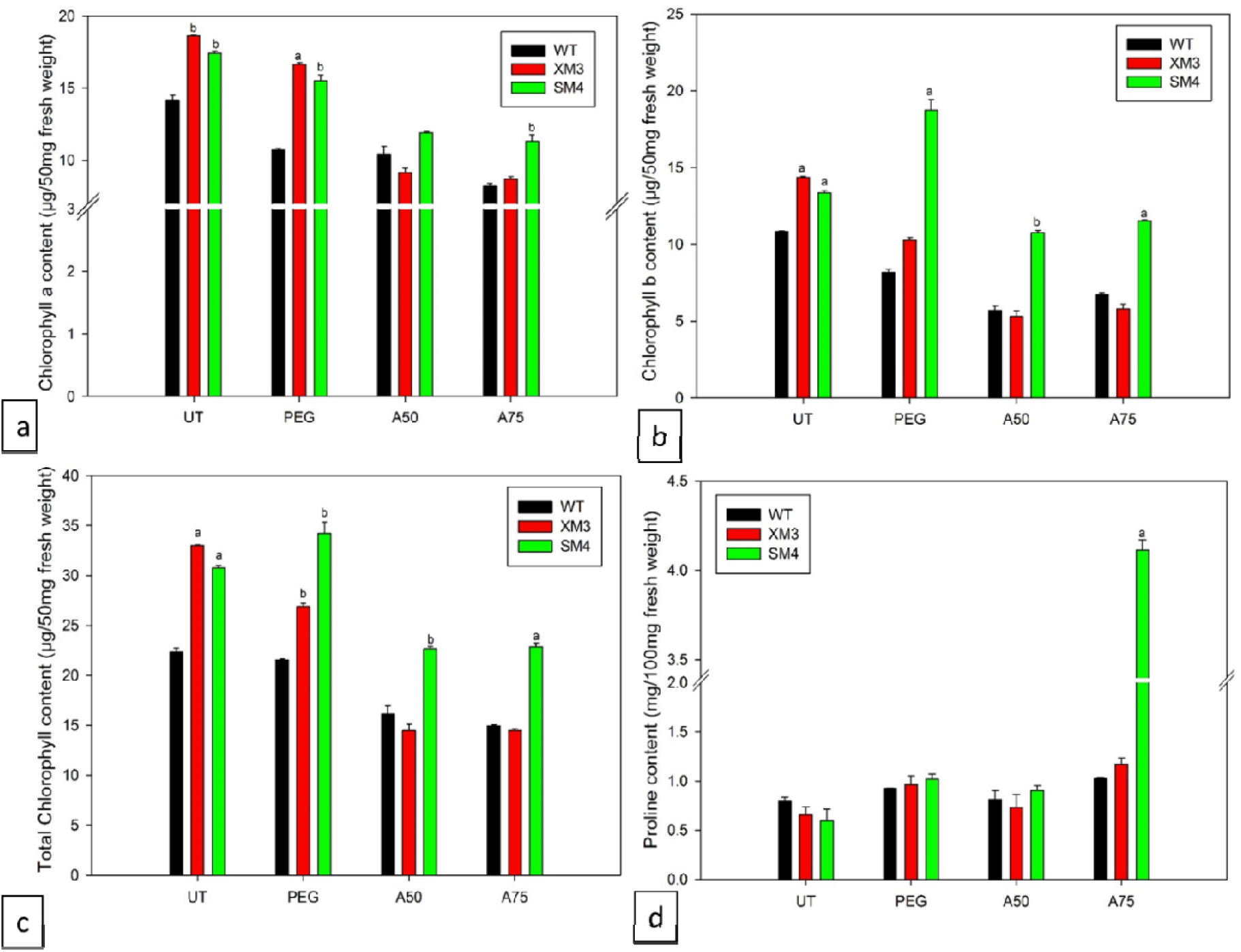
Biochemical analysis of the tagged lines. Graphical representation of the biochemical studies done on XM3 and SM4 in comparison to WT plants post 20 days of stress. (a), (b), (c), (d) depicts chlorophyll a, chlorophyll b, total chlorophyll and proline content post stress respectively. One way ANOVA was performed at a significance level P<0.001 marked as a, P<0.025 marked as b, P<0.005 marked as c.

**Figure 5:**
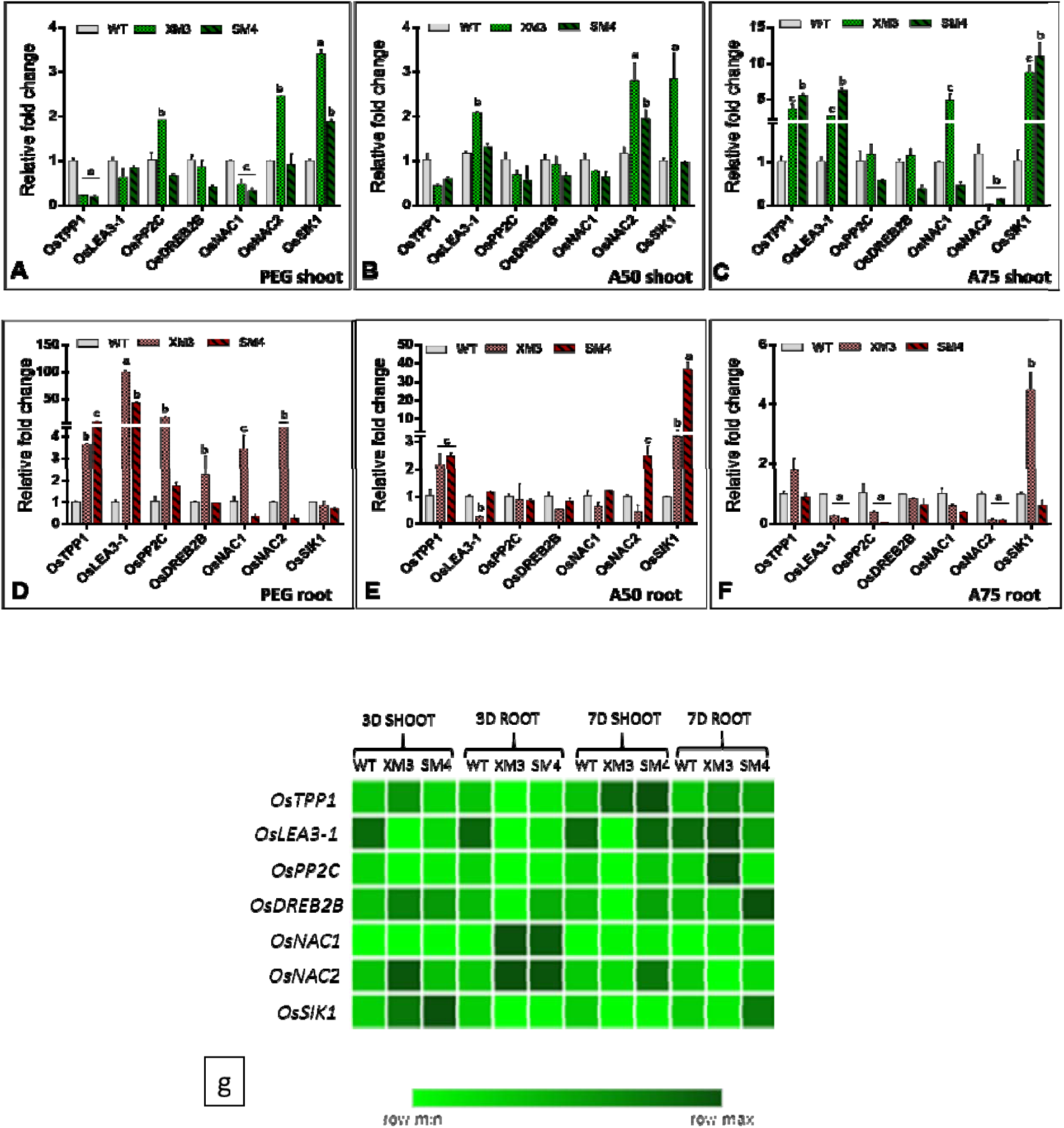
Expression analysis of stress responsive genes under simulated and pot-level drought conditions. Graph representing the transcript level of seven drought specific genes under imposed stress cues (10% PEG, ABA 50µM and ABA 75µM) in shoot (A to C) and root (D to F) tissues of the tagged lines with respect to the WT plants. Rice *actin* was used as the internal reference gene. The individual WT sample for each treatment was used to normalise the data. The fold change was calculated using ΔΔC_T_ method. The mean and the standard error is plotted in a vertical bar graph. One way ANOVA was performed at a significance level P<0.001 marked as a, P<0.025 marked as b and P<0.05 marked as c. (F) Transcript analysis of seven drought specific genes in root and shoot tissues post 3days and 7 days drought. The data was normalized using rice *actin* as the internal reference gene. The corresponding WT samples for each drought treatment was used to normalise the data, and the fold change was calculated using ΔΔC_T_ method. The results are depicted in the form of a heat map generated by the MORPHEUS program.

In SM4, the chlorophyll a, b and total chlorophyll contents post stress ranged from 11 to 15 µg, 11 to 19 µg and 23 to 34 µg/50mg fresh weight while that of the WT ranged from 8 to 11 µg, 5 to 8 µg and 15 to 21 µg/50 mg fresh weight, respectively.

Under untreated conditions, WT, XM3 and SM4 had similar proline content which ranged from 0.6 to 0.7 mg/100 mg fresh weight. Treatment with 10% PEG and 50 µM ABA had induced the proline content to increase up to 0.9 to 1.0mg/100mg fresh weight in all of them. Under ABA 75µM treatment, SM4 accumulated a very high proline content (4 mg/100 mg fresh weight) compared with WT (Fig. 4d). After revival, the proline content of all the treated lines dropped and was almost similar to their corresponding untreated controls (Fig. S7d). Thus, higher chlorophyll and proline contents might have resulted in better photosynthetic efficiency and drought stress tolerance mechanism of the tagged lines leading to a sustainable productivity.

### 3.6. Yield-related observations of the tagged lines under pot-level drought conditions

The tagged lines and the WT plants were subjected to periodic drought condition in pots for three (60% FC) and seven (40% FC) days. The WT showed lesser green phenotype and more leaf rolling compared to XM3 and SM4 after 7 days (Fig. S8). No significant difference in terms of yield and phenotypic parameters was observed post 3 days of drought. After induction of consecutive drought for 7 days, WT yielded ∼200 seeds per plant and each of two mutants produced ∼400 seeds per plant. Accordingly, the number of primary branches per panicle, number of seeds per primary branch and the number of seeds per panicle were also decreased in WT. The mean and the standard error of the data collected are provided in the Table 1.

### 3.7. Seedling germination assay

The seed germination in 50 and 75 µM ABA showed a germination retardation after 5 days compared with the untreated control (Fig. S9). The seeds of the tagged mutant lines continued to germinate and survived on ABA containing medium. Both XM3 and SM4 had longer roots with emerging shoots even at higher concentrations of ABA whereas, the WT seeds showed mild germination under 50 µM concentration and completely failed to germinate at 75 µM. These results might suggest that higher expression of SEN1 and XPB2 renders the plant less sensitivity to ABA.

### 3.8. Transcriptional analysis of stress responsive genes in tagged lines

We checked the transcript levels of seven stress responsive genes (*OsTPP1, OsLEA3-1, OsPP2C, OsDREB2B, OsNAC1, OsNAC2* and *OsSIK1)* in the shoot (Fig. 6a to c) and root (Fig. 6d to f) tissues of tagged lines under treated conditions (PEG, 50 and 75 µM ABA) with respect to corresponding WT. Most of these genes in shoot were upregulated under the influence of ABA treatment, while several of them were upregulated under PEG in roots. This showed that the helicases might have an influence on the expression of certain stress responsive genes resulting in stress tolerance. Two genes, *OsPP2C* (ABA-dependent) and *OsDREB2B* (ABA-independent) was observed to be either downregulated or have equal level of expression as the WT in shoot and root tissues under ABA. The observation was constant in both XM3 and SM4 tagged lines. This might be an indication that XPB2 and SEN1 are involved in both ABA dependent and independent pathways of stress tolerance.

In shoot tissues of XM3 lines, ABA 50µM induced the expression of *OsNAC1*, and, *OsTPP1* and *OsNAC1* were upregulated by 3 to 8 fold under ABA 75µM treatment. *OsLEA3-1* and *OsSIK1* was induced (2 to 9 fold) under both concentrations of ABA. Under PEG treatment, *OsSIK1, OsNAC2* and *OsPP2C* showed 2-3 fold upregulation. In roots, *OsTPP1, OsLEA3-1, OSPP2C, OsDREB2B, OsNAC1* and *OsNAC2* were upregulated all under PEG. *OsLEA3-1* and *OsPP2C* exhibited highest transcript levels up to 100 fold and 15 fold, respectively. In response to both 50 and 75 µM ABA, *OsTPP1* and *OsSIK1* showed 2-4 fold upregulation in XM3 lines.

The shoots of SM4 showed 5 to 6 fold upregulation of *OsTPP1, OsLEA3-1* under 75µM ABA and 2 fold upregulation of *OsNAC*2 under 50 µM ABA.. *OsSIK1* was highly upregulated by 8 fold and 11 fold under PEG and ABA 75µMtreatment, respectively. In root tissues, *OsTPP1, OsLEA3-1, OSPP2C* were upregulated under PEG treatment and *OsLEA3-1* exhibited the highest transcript level up to 42 fold. Under ABA 50µM treatment, *OsTPP1* and *OsNAC2* was expressed by 2.5 fold and *OsSIK1* was upregulated by 37 fold in SM4 lines. *OsTPP1* and *OsSIK1* was upregulated under both concentrations of ABA.

We have also performed similar analysis on the tagged lines after subjecting them to pot-level drought stress for 3 and 7 days (Fig. 6g). It was observed that the upregulation of these genes was more prominent in the root tissues when compared to the shoots. Under pot-level drought conditions, *OsNAC2* and *OsSIK1* showed a moderate upregulation of 2-3 folds in shoots after 3 days, whereas in roots, *OsNAC1* and *OsNAC2* were upregulated up to 7 fold and 2 fold, respectively in XM3 lines. In SM4 lines, *OsSIK1* showed 3.5 fold upregulation in shoot and *OsNAC1* and *OsNAC2* showed 7 and 2 fold upregulation in root. Thus, after 3 days, a total of three out of seven genes under study were upregulated. After 7 days of continuous drought treatment, six out of seven genes became upregulated. 2 fold upregulation of *OsTPP1* was observed in XM3 shoots, whereas, *OsPP2C* showed 4 fold upregulation in roots. In SM4 shoots, *OsTPP1* and *OsNAC2* genes showed a 2 fold upregulation and *OsDREB2B, OsNAC1* and *OsSIK1* were upregulated by 2.6 fold in SM4 roots.

## 4. Discussion

The functions of redundant and lethal genes in plants can be effectively studied by integrating tetrameric enhancers that activate the surrounding genes in either direction in plant genome by a mechanism called activation tagging (Weigel et al., 2000; Jeong et al., 2002; Wan et al., 2009). Some of the activation tagged lines generated earlier by our group showed high WUE-related physiological parameters and yield under water limited conditions (Moin et al, 2016). The individual CaMV35S promoters cause ectopic overexpression of gene(s) that are in transcriptional fusion with them which sometimes lead to pleiotropic effects, whereas the enhancers which are modified versions of 35S promoters do not lead to constitutive expression but upregulate the target genes more than their endogenous levels. Thus, the plant phenotype resulting from enhancer-mediated activation would more likely reflect the normal function of the tagged gene (Weigel et al. 2000; Jeong et al., 2002; Tani et al. 2002).

### 4.1 *XPB2* and *SEN1* helicases are induced in rice roots under chemical and hormonal treatments

The two mutants used in this study were stable (*Ac*^-^/*Ds*^+^) with the absence of *Ac* element. Also, the enhancer integration was intergenic and hence, the function of other endogenous genes might not have disrupted which would have otherwise occurred as a result of intragenic inactivation. Because of stable and intergenic integration, the WUE and drought tolerant phenotypes of the two selected mutants have resulted more likely by enhanced activation of their respective genes. The mutant lines were shortlisted based on their quantum efficiency and Δ^13^C values. During water stress, plants tend to close their stomatal aperture, thereby reducing their intercellular CO_2_ concentration (C_i_). Δ^13^C is directly influenced by the C_i_ and hence its decrease during stress increases the carbon discrimination and lowers the WUE (Martin & Thorstenson, 1988; Chen et al., 2011). The higher quantum efficiency and low Δ^13^C of mutants as observed in this study indicates high photosynthetic performance and WUE, respectively, thereby resulted in increased yield under water withdrawal conditions. In XM3 line, *XPB2* DNA helicase, situated next to the enhancer was upregulated as compared to other genes present in the vicinity, which showed no upregulation as WT. In a similar way, *SEN1* RNA helicase in SM4 located in vicinity to the enhancer became upregulated.

The expression of *SEN1* and *XPB2* was induced particularly in root tissues by external application of stress inducible phytohormone ABA and drought inducing agent, PEG. Since roots are the first organs to perceive stress, an upregulation of the genes in these tissue indicates their possible role in the regulation of a signaling cascade for drought stress responses (Janiak et al., 2015). Unlike abiotic stress treatments, the transcript levels of *SEN1* and *XPB2* were not significantly induced by *Xoo* and *R. solani* pathogens.

### 4.2 XPB2 and SEN1 help in maintaining sustainable productivity in rice under limited water conditions

The SEN1 and XPB2 gain of function mutants were subjected to drought experiments by allowing them to grow in PEG and two concentrations of ABA solution and water withdrawal conditions in pots. In all the sets of experiments, the tagged lines, XM3 and SM4 performed well under the stress conditions. Root length under water deficiency is an important trait for plant productivity. Thus, comparatively longer roots in SM4 lines under simulated stress is an indication of its enhanced drought tolerance (Janiak et al., 2015; Sharp and LeNoble, 2002). Similarly, the tagged lines showed healthy phenotypes after 3 and 7 days of drought treatment in pots with respect to the WT. Drought hampers the water balance of the plant, causing growth retardation and early senescence of leaves (Anjum et al., 2011; Sreenivasulu et al., 2012). We observed similar phenomena in the WT, which had a lower biomass, higher wilting percentage and lower plant revival percentage compared with the tagged mutant counterparts under simulated stress conditions. Therefore, SEN1 and XPB2 appeared to be involved in maintaining the rigidity of the plant during and after stresses.

High level of ABA leads to the inhibition of shoot growth, less carbon accumulation and yield (Zhang et al., 2006; Blum, 2005). Such stress avoidance mechanism was found in the WT, which showed decreased plant height and yield post revival. The exogenous stress cues had little impact on the physiology and yield of the gain of function mutant lines. Prolonged drought closes the stomata causing ineffective gas exchange, and a decrease in the photosynthetic capability of the plant (Sreenivasulu et al., 2012). This is evident from the photosynthetic performance of the WT in comparison to XM3 and SM4 lines. The Fv/Fm ratio as measured by PAM (Pulse Amplitude Modulated) fluorometer indicates the photochemical efficiency or the level of stress as experienced by the plant (Murchie and Lawson, 2013; Osmolovskaya et al., 2018). The healthy untreated WT and mutant plants had a quantum efficiency ranging from 74 to 77%. When exposed to stress, the efficiency of the WT decreased to a range of 63-67%, while that of the tagged lines remained nearly unaltered (72-77%). This is indicative of the fact that the photosynthetic ability of the tagged lines was not deterred upon exposure to stress. Since both chlorophyll a and b are susceptible towards drought stress, their higher content in the tagged lines corroborates better photosynthetic capability of XM3 and SM4 and, in turn, their yield (Anjum et al., 2011; Osmolovskaya et al., 2018; Zhang et al., 2018). Similarly, SM4 lines also accumulated a high level of proline under ABA treatment. During drought stress, plants maintain turgor pressure by accumulating ions and organic solutes and proline is one of these compatible solutes. Higher level of this constitutes a known osmotic adjustment in cells. Greater accumulation of osmoprotectants indicate better drought tolerance and regulation of osmotic potential (Anjum et al., 2011; Yang et al., 2018; Zhang et al., 2018).

The boot leaves are considered as an important source of metabolites in rice plants that contribute to plant productivity. Their length has a positive correlation with the panicle length and yield (Rahman et al., 2013). Also, high ABA accumulation in reproductive tissues can lead to abortion of pods and reduce grain filling period (Sreenivasulu et al., 2012). Therefore, a decrease in the boot leaf and panicle length in the WT under stress is an indication of its susceptibility towards drought. A 26-98% more yield per plant in the tagged lines over the WT under simulated conditions and 9-50% more yield per tagged line under pot-level drought experiments implies that the SEN1 and XPB2 helicases are responsible for maintaining a sustainable yield in the gain-of-function mutants even under severe stress conditions. The helicases identified in the present study do not belong to the DEAD/H box helicases, which are well-known for imparting drought tolerance when overexpressed in plants. These include OsSUV3 from rice whose overexpression in IR64 rice variety has led to improved drought tolerance (Tuteja et al., 2013). Another rice RNA helicase OsRH58 exhibited drought tolerant activity in Arabidopsis (Nawax & Kang, 2019). PDH45 and PDH47 are reported pea DNA helicases, which have similar tolerance when expressed under the constitutive 35S promoter in groundnut (Manjulatha et al., 2014), chili (Shivakumara et al., 2017) and rice (Singha et al., 2017). SIDEAD31, a RNA helicase from Arabidopsis, imparted drought tolerance when expressed constitutively in tomato. (Zhu et al., 2015). Other helicases from Arabidopsis include RH8, RH9 and RH25, which had been reported for enhancing drought tolerance in plants (Kant et al., 2007; Baek et al., 2018).

The WT seeds inoculated on 75 µM concentration of ABA failed to germinate after 5 days, but XM3 and SM4 tagged lines germinated normally and continued to grow. Usually high ABA concentration compromises the germination rate and seedling establishment causing cell dehydration, wilting and death. (Zhang et al., 2018). Germination ability of tagged lines indicates the probable role of SEN1 and XPB2 in negative regulation of ABA mediated seed dormancy.

### 4.3 Existence of a probable cross-talk between SEN1 and XPB2 with other stress regulatory pathways resulting in drought tolerance of rice

The expression level of seven stress-specific genes was studied in the tagged lines under simulated drought stress (PEG and ABA) and pot level drought after the complete withdrawal of water. There was an upregulation of all seven genes in XM3 and five genes in SM4 under simulated stress conditions. Under pot-level drought, four genes in XM3 and seven genes in SM4 became upregulated. This indicates a possible cross-talk between *SEN1*/*XPB2* and other stress-tolerant genes, which together not only induced stress-tolerance in the tagged lines but also maintained their productivity under stress. However, the interaction between the stress regulatory genes and the pathways they operate requires further investigation.

In conclusion, the gain of function mutant lines of *SEN1* and *XPB2* genes had the advantage over the WT in overcoming drought stress. Currently, efforts are underway to independently overexpress these two genes for a detailed functional characterization and also to understand the underlying mechanism of stress tolerance. The expression of these genes is regulated by ABA, PEG and other stress factors, but there is no conclusive evidence that these genes work in an ABA-dependant or an independent manner. The promoter analysis of *XPB2* showed the existence of ABRE, DRE and MYB *cis*-acting elements and that of *SEN1* showed the presence of MYB and ABRE motifs. Presence of both DRE and ABRE elements suggests the simultaneous existence of ABA dependent and independent gene regulation. (Roychoudhury et al, 2013; Yosida et al., 2014) Such phenomenon had been noticed in Arabidopsis *rd29A* gene promoter (Narusaka et al. 2003). Also, MYB transcription factors are mostly known to work in an ABA dependent manner during stress, but there is evidence where certain MYB factor like OsMYB3R-2 was shown to be less sensitive to ABA (Dai et al., 2007). From our observation it became clear that the XM3 and SM4 gain-of-function mutant lines have decreased sensitivity towards ABA although their expression is induced by it. Thus SEN1 and XPB2 helicases might be playing an intermediate role in ABA dependent and independent pathway. *OsPP2C* is a very potent ABA dependent gene and *OsDREB2B* is another ABA independent gene either shows equal level of expression as the control or gets downregulated both in root and shoot. The downregulation or unchanged expression of *OsPP2C* (an ABA-dependent gene) and *OsDREB2B* (an ABA-independent gene) during the expression analysis under ABA treatment is probably an outcome of crosstalk between the two pathways or the existence of a negative feedback mechanism. Reports say the existence of such negative regulation of stress marker genes by *OsNAC2* inspite of the gene itself getting induced by ABA. (Shen at al., 2017). Based on the previous studies, it can also be assumed that both genes regulate stress tolerance mechanism either by inducing DNA repair pathways to overcome DNA damage resulting from stress or by effectively terminating pervasive transcription, or by resolving unwanted DNA: RNA / RNA: RNA hybrids formed during stress (Mischo et al., 2011; Richards et al., 2008; Raikwar et al., 2015; Han et al., 2017; Mischo et al., 2018) and thereby, upregulating the transcription of other important stress regulatory genes. These genes appear to have a positive regulation towards stress tolerance of rice plants.

## Author contribution statement

PBK, MM and MD designed the experiments and prepared the manuscript. MD performed the experiments. MM and AB generated and maintained the activation tagged mutant lines. AS helped in the qRT-PCR experiments, analysis and seed counting. PBK supervised the work. MD, MM and PBK prepared the manuscript.

## Acknowledgements

MD is grateful to UoH BBL and UGC for research fellowship and to DBT for funding the rice activation tagging project. MM is thankful to DST for fellowship and research grant in the form of INSPIRE-faculty program. AS and AB are grateful to DBT for proving Research Fellowships. MD is grateful to Dr. M.S. Madhav, Department of Biotechnology, Indian Institute of Rice Research, Hyderabad, India, for providing the infected rice samples and wild type BPT-5204 seeds. PBK is grateful to the National Academy of Sciences-India for the NASI-Platinum Jubilee Senior Scientist award. The authors are grateful to Prof. M.Udaya Kumar, Bengaluru, India for his help in IRMS analysis.

## Abbreviations

SEN1: t-RNA Splicing Endonuclease
XPB2: Xeroderma Pigementosa group B2
WUE: water use efficiency
PEG: Polyethylene Glycol
ABA: Abscisic acid

## References

Ambawat, S., Sharma, P., Yadav, N. R., & Yadav, R. C. (2013). MYB transcription factor genes as regulators for plant responses: An overview. Physiology and Molecular Biology of Plants, 19(3), 307–321. https://doi.org/10.1007/s12298-013-0179-1

Anjum, S. A., Xie, X. yu, Wang, L. chang, Saleem, M. F., Man, C., & Lei, W. (2011). Morphological, physiological and biochemical responses of plants to drought stress. African Journal of Agricultural Research, 6(9), 2026–2032. https://doi.org/10.5897/AJAR10.027

Arciga-Reyes, L., Wootton, L., Kieffer, M., & Davies, B. (2006). UPF1 is required for nonsense-mediated mRNA decay (NMD) and RNAi in Arabidopsis. Plant Journal, 47(3), 480–489. https://doi.org/10.1111/j.1365-313X.2006.02802.x

Baek, W. (2018). *A DEAD* - *box RNA helicase, RH8, is critical for regulation of ABA signalling and the drought stress response via inhibition of PP2CA activity*. March, 1593–1604. https://doi.org/10.1111/pce.13200

Bhatia, Prakash K., Wang, Z., & Friedberg, E. C. (1996). DNA repair and transcription. Current Opinion in Genetics and Development. https://doi.org/10.1016/S0959-437X(96)80043-8

Blum, A. (2005). Drought resistance, water-use efficiency, and yield potential - Are they compatible, dissonant, or mutually exclusive? Australian Journal of Agricultural Research. https://doi.org/10.1071/AR05069

Caine, R. S., Yin, X., Sloan, J., Harrison, E. L., Mohammed, U., Biswal, A. K., Dionora, J., Chater, C. C., Coe, R. A., Bandyopadhyay, A., Murchie, E. H., Swarup, R., Quick, W. P., Gray, J. E., & Gray, J. E. (2019). Rice with reduced stomatal density conserves water and has improved drought tolerance under future climate conditions. 371–384. https://doi.org/10.1111/nph.15344

Çakir, B., Kiliçkaya, O., & Olcay, A. C. (2013). Genome-wide analysis of Aux/IAA genes in Vitis vinifera: Cloning and expression profiling of a grape Aux/IAA gene in response to phytohormone and abiotic stresses. Acta Physiologiae Plantarum, 35(2), 365–377. https://doi.org/10.1007/s11738-012-1079-7

Chabouté, M. E., Clément, B., & Philipps, G. (2002). S phase and meristem-specific expression of the tobacco RNR1b gene is mediated by an E2F element located in the 5′ leader sequence. Journal of Biological Chemistry, 277(20), 17845–17851. https://doi.org/10.1074/jbc.M200959200

Costa, R. M. A., Morgante, P. G., Berra, C. M., Nakabashi, M., Bruneau, D., Bouchez, D., Sweder, K. S., Van Sluys, M. A., & Menck, C. F. M. (2001). The participation of AtXPB1, the XPB/RAD25 homologue gene from Arabidopsis thaliana, in DNA repair and plant development. Plant Journal, 28(4), 385–395. https://doi.org/10.1046/j.1365-313X.2001.01162.x

Dai, X., Xu, Y., Ma, Q., Xu, W., Wang, T., Xue, Y., & Chong, K. (2007). Overexpression of an R1R2R3 MYB gene, OsMYB3R-2, increases tolerance to freezing, drought, and salt stress in transgenic Arabidopsis. Plant Physiology, 143(4), 1739–1751. https://doi.org/10.1104/pp.106.094532

Ding, S., He, F., Tang, W., Du, H., & Wang, H. (2019). Identification of maize cc-type glutaredoxins that are associated with response to drought stress. Genes, 10(8). https://doi.org/10.3390/genes10080610

Guzder, S. N., Habraken, Y., Sung, P., Prakash, L., & Prakash, S. (1995). Reconstitution of yeast nucleotide excision repair with purified Rad proteins, replication protein A, and transcription factor TFIIH. Journal of Biological Chemistry. https://doi.org/10.1074/jbc.270.22.12973

Haefele, S. M., Kato, Y., & Singh, S. (2016). Field Crops Research Climate ready rice[: Augmenting drought tolerance with best management practices. Field Crops Research, 190, 60–69. https://doi.org/10.1016/j.fcr.2016.02.001

Han, Z., Libri, D., & Porrua, O. (2017). Biochemical characterization of the helicase Sen1 provides new insights into the mechanisms of non-coding transcription termination. Nucleic Acids Research, 45(3), 1355–1370. https://doi.org/10.1093/nar/gkw1230

Janiak, A., Kwasniewski, M., & Szarejko, I. (2015). Gene expression regulation in roots under drought. December. https://doi.org/10.1093/jxb/erv512

Jankowsky, E., & Fairman, M. E. (2007). RNA helicases - one fold for many functions. Current Opinion in Structural Biology, 17(3), 316–324. https://doi.org/10.1016/j.sbi.2007.05.007

Jeong, D.-H., An, S., Kang, H.-G., Moon, S., Han, J.-J., Park, S., Lee, H. S., An, K., & An, G. (2002). T-DNA Insertional Mutagenesis for Activation Tagging in Rice. Plant Physiology, 130(December), 1636–1644. https://doi.org/10.1104/pp.014357.

Jeong, D. H., An, S., Kang, H. G., Moon, S., Han, J. J., Park, S., Sook Lee, H., An, K., & An, G. (2002). T-DNA insertional mutagenesis for activation tagging in rice. Plant Physiology. https://doi.org/10.1104/pp.014357

Leonaitė, B., Han, Z., Basquin, J., Bonneau, F., Libri, D., Porrua, O., & Conti, E. (2017). Sen1 has unique structural features grafted on the architecture of the Upf1-like helicase family. The EMBO Journal, 36(11), 1590–1604. https://doi.org/10.15252/embj.201696174

Lescot, M., Déhais, P., Thijs, G., Marchal, K., Moreau, Y., Van de Peer, Y., & Rombauts, S. (2002). PlantCARE, a database of plant cis-acting regulatory elements and a portal to tools for in silico analysis of promoter sequences. Nucleic acids research, 30(1), 325–327.

Li, W., Selvam, K., Rahman, S. A., & Li, S. (2016). Sen1, the yeast homolog of human senataxin, plays a more direct role than Rad26 in transcription coupled DNA repair. Nucleic Acids Research. https://doi.org/10.1093/nar/gkw428

Linder, P., & Owttrim, G. W. (2009). Plant RNA helicases: linking aberrant and silencing RNA. Trends in Plant Science, 14(6), 344–352. https://doi.org/10.1016/j.tplants.2009.03.007

Lo, S., Fan, M., Hsing, Y., Chen, L., Chen, S., Wen, I., Liu, Y., & Chen, K. (2020). Genetic resources offer efficient tools for rice functional genomics research. 2016, 998–1013. https://doi.org/10.1111/pce.12632

Luo, Y., Liu, Y. B., Dong, Y. X., Gao, X., & Ã, X. S. Z. (2009). Expression of a putative alfalfa helicase increases tolerance to abiotic stress in Arabidopsis by enhancing the capacities for ROS scavenging and osmotic adjustment. 166. https://doi.org/10.1016/j.jplph.2008.06.018

Manjulatha, M., Sreevathsa, R., Kumar, A. M., Sudhakar, C., Prasad, T. G., Tuteja, N., & Udayakumar, M. (2014). Overexpression of a Pea DNA Helicase (PDH45) in Peanut (Arachis hypogaea L .) Confers Improvement of Cellular Level Tolerance and Productivity Under Drought Stress. 111–125. https://doi.org/10.1007/s12033-013-9687-z

Martin-Tumasz, S., & Brow, D. A. (2015). Saccharomyces cerevisiae sen1 helicase domain exhibits 5′-to 3′-helicase activity with a preference for translocation on DNA rather than RNA. Journal of Biological Chemistry, 290(38), 22880–22889. https://doi.org/10.1074/jbc.M115.674002

Mischo, H. E., Chun, Y., Harlen, K. M., Smalec, B. M., Dhir, S., Churchman, L. S., & Buratowski, S. (2018). Cell-Cycle Modulation of Transcription Termination Factor Sen1. Molecular Cell, 70(2), 312-326.e7. https://doi.org/10.1016/j.molcel.2018.03.010

Mischo, H. E., Gómez-González, B., Grzechnik, P., Rondón, A. G., Wei, W., Steinmetz, L., Aguilera, A., & Proudfoot, N. J. (2011). Yeast Sen1 helicase protects the genome from transcription-associated instability. Molecular Cell, 41(1), 21–32. https://doi.org/10.1016/j.molcel.2010.12.007

Moin, M., Bakshi, A., Saha, A., Udaya Kumar, M., Reddy, A. R., Rao, K. V., Siddiq, E. A., & Kirti, P. B. (2016). Activation tagging in indica rice identifies ribosomal proteins as potential targets for manipulation of water-use efficiency and abiotic stress tolerance in plants. Plant Cell and Environment, 39(11), 2440–2459. https://doi.org/10.1111/pce.12796

Moin, M., Bakshi, A., Madhav, M. S., & Kirti, P. B. (2017). Expression profiling of ribosomal protein gene family in dehydration stress responses and characterization of transgenic rice plants overexpressing RPL23A for water-use efficiency and tolerance to drought and salt stresses. Frontiers in chemistry, 5, 97.

Morgante, P. G., Berra, C. M., Nakabashi, M., Costa, R. M. A., Menck, C. F. M., & Van Sluys, M. A. (2005). Functional XPB/RAD25 redundancy in Arabidopsis genome: Characterization of AtXPB2 and expression analysis. Gene, 344, 93–103. https://doi.org/10.1016/j.gene.2004.10.006

Murchie, E. H., & Lawson, T. (2013). Chlorophyll fluorescence analysis: A guide to good practice and understanding some new applications. Journal of Experimental Botany, 64(13), 3983–3998. https://doi.org/10.1093/jxb/ert208

Narusaka, Y., Nakashima, K., Shinwari, Z. K., Sakuma, Y., Furihata, T., Abe, H., Narusaka, M., Shinozaki, K., & Yamaguchi-Shinozaki, K. (2003). Interaction between two cis-acting elements, ABRE and DRE, in ABA-dependent expression of Arabidopsis rd29A gene in response to dehydration and high-salinity stresses. Plant Journal, 34(2), 137–148. https://doi.org/10.1046/j.1365-313X.2003.01708.x

Nawaz, G., & Kang, H. (2019). Rice OsRH58, a chloroplast DEAD-box RNA helicase, improves salt or drought stress tolerance in Arabidopsis by affecting chloroplast translation. 1–11.

Osmolovskaya, N., Shumilina, J., Kim, A., Didio, A., Grishina, T., Bilova, T., Keltsieva, O. A., Zhukov, V., Tikhonovich, I., Tarakhovskaya, E., & Frolov, A. (2018). Methodology of Drought Stress Research□: Experimental Setup and Physiological Characterization. https://doi.org/10.3390/ijms19124089

Park, E., Guzder, S. N., Koken, M. H. M., Jaspers-Dekker, I., Weeda, G., Hoeijmakers, J. H. J., Prakash, S., & Prakash, L. (1992). RAD25 (SSL2), the yeast homolog of the human xeroderma pigmentosum group B DNA repair gene, is essential for viability. Proceedings of the National Academy of Sciences of the United States of America, 89(23), 11416–11420. https://doi.org/10.1073/pnas.89.23.11416

Passricha, N., Saifi, S. K., Gill, S. S., Tuteja, R., & Tuteja, N. (2018). Role of Plant Helicases in Imparting Salinity Stress Tolerance to Plants. In Helicases from All Domains of Life (Issue 3). Elsevier Inc. https://doi.org/10.1016/B978-0-12-814685-9.00003-8

Qu, S., Desai, A., Wing, R., & Sundaresan, V. (2008). A versatile transposon-based activation tag vector system for functional genomics in cereals and other monocot plants. Plant Physiology, 146(1), 189–199. https://doi.org/10.1104/pp.107.111427

Rahman, M. A., Haque, M., Sikdar, B., Islam, M. A., & Matin, M. N. (2014). Correlation Analysis of Flag Leaf with Yield in Several Rice Cultivars. Journal of Life and Earth Science, 8, 49–54. https://doi.org/10.3329/jles.v8i0.20139

Raikwar, S., Srivastava, V. K., Gill, S. S., Tuteja, R., & Tuteja, N. (2015). Emerging importance of helicases in plant stress tolerance: Characterization of oryza sativa repair helicase XPB2 promoter and its functional validation in tobacco under multiple stresses. Frontiers in Plant Science, 6(DEC), 1–7. https://doi.org/10.3389/fpls.2015.01094

Richards, J. D., Cubeddu, L., Roberts, J., Liu, H., & White, M. F. (2008). The Archaeal XPB Protein is a ssDNA-Dependent ATPase with a Novel Partner. Journal of Molecular Biology, 376(3), 634–644. https://doi.org/10.1016/j.jmb.2007.12.019

Robin, A. H. K., Uddin, M. J., & Bayazid, K. N. (2015). Polyethylene glycol (PEG)-treated hydroponic culture reduces length and diameter of root hairs of wheat varieties. Agronomy, 5(4), 506–518. https://doi.org/10.3390/agronomy5040506

Roychoudhury, A., Paul, S., & Basu, S. (2013). Cross-talk between abscisic acid-dependent and abscisic acid-independent pathways during abiotic stress. In Plant Cell Reports (Vol. 32, Issue 7, pp. 985–1006). https://doi.org/10.1007/s00299-013-1414-5

Sakai, T., Takahashi, Y., & Nagata, T. (1996). Analysis of the promoter of the auxin-inducible gene, parC, of tobacco. Plant and Cell Physiology, 37(7), 906–913. https://doi.org/10.1093/oxfordjournals.pcp.a029038

Sariki, S. K., Sahu, P. K., Golla, U., Singh, V., Azad, G. K., & Tomar, R. S. (2016). Sen1, the homolog of human Senataxin, is critical for cell survival through regulation of redox homeostasis, mitochondrial function, and the TOR pathway in Saccharomyces cerevisiae. FEBS Journal, 283(22), 4056–4083. https://doi.org/10.1111/febs.13917

Seraj, Z. I., Elias, S. M., Biswas, S., & Tuteja, N. (2018). Helicases and Their Importance in Abiotic Stresses. In Salinity Responses and Tolerance in Plants (Vol. 2). https://doi.org/10.1007/978-3-319-90318-7

Sharp, R. E., & LeNoble, M. E. (2002). ABA, ethylene and the control of shoot and root growth under water stress. Journal of Experimental Botany. https://doi.org/10.1093/jexbot/53.366.33

Shen, J., Lv, B., Luo, L., He, J., Mao, C., Xi, D., & Ming, F. (2017). The NAC-type transcription factor OsNAC2 regulates ABA-dependent genes and abiotic stress tolerance in rice. June 2016, 1–14. https://doi.org/10.1038/srep40641

Shivakumara, T. N., Sreevathsa, R., Dash, P. K., & Sheshshayee, M. S. (2017). Overexpression of Pea DNA Helicase 45 (PDH45) imparts tolerance to multiple abiotic stresses in chili (Capsicum annuum L .). April, 1–12. https://doi.org/10.1038/s41598-017-02589-0

Singha, D. L., Tuteja, N., & Boro, D. (2017). Heterologous expression of PDH47 confers drought tolerance in indica rice. Plant Cell, Tissue and Organ Culture (PCTOC), 130(3), 577–589. https://doi.org/10.1007/s11240-017-1248-x

Sloan, K. E., & Bohnsack, M. T. (2018). Unravelling the Mechanisms of RNA Helicase Regulation. Trends in Biochemical Sciences, 43(4), 237–250. https://doi.org/10.1016/j.tibs.2018.02.001

Sreenivasulu, N., Harshavardhan, V. T., Govind, G., Seiler, C., & Kohli, A. (2012). Contrapuntal role of ABA: Does it mediate stress tolerance or plant growth retardation under long-term drought stress? Gene, 506(2), 265–273. https://doi.org/10.1016/j.gene.2012.06.076

Srivastav, A., Mehta, S., Lindlof, A., & Bhargava, S. (2010). Over-represented promoter motifs in abiotic stress-induced DREB genes of rice and sorghum and their probable role in regulation of gene expression. Plant Signaling and Behavior, 5(7), 775–784. https://doi.org/10.4161/psb.5.7.11769

Steinmetz, E. J., Warren, C. L., Kuehner, J. N., Panbehi, B., Ansari, A. Z., & Brow, D. A. (2006). Genome-Wide Distribution of Yeast RNA Polymerase II and Its Control by Sen1 Helicase. Molecular Cell, 24(5), 735–746. https://doi.org/10.1016/j.molcel.2006.10.023

Tani, H., Chen, X., Nurmberg, P., Grant, J. J., SantaMaria, M., Chini, A., … & Loake, G. J. (2004). Activation tagging in plants: a tool for gene discovery. Functional & integrative genomics, 4(4), 258–266.

Tuteja, N. (2003). Plant DNA helicases: The long unwinding road. Journal of Experimental Botany, 54(391), 2201–2214. https://doi.org/10.1093/jxb/erg246

Tuteja, N., Sahoo, R. K., Garg, B., Tuteja, R., Asaf, A., & Marg, A. (2013). OsSUV3 dual helicase functions in salinity stress tolerance by maintaining photosynthesis and antioxidant machinery in. 115–127. https://doi.org/10.1111/tpj.12277

Umate, P., Tuteja, R., & Tuteja, N. (2010). Genome-wide analysis of helicase gene family from rice and Arabidopsis: A comparison with yeast and human. Plant Molecular Biology, 73(4), 449–465. https://doi.org/10.1007/s11103-010-9632-5

Wan, S., Wu, J., Zhang, Z., Sun, X., Lv, Y., Gao, C., Ning, Y., Ma, J., Guo, Y., Zhang, Q., Zheng, X., Zhang, C., Ma, Z., & Lu, T. (2009). Activation tagging, an efficient tool for functional analysis of the rice genome. Plant Molecular Biology, 69(1–2), 69–80. https://doi.org/10.1007/s11103-008-9406-5

Wei, Q., Cao, H., Li, Z., Kuai, B., & Ding, Y. (2013). Identification of an AtCRN1-like chloroplast protein BeCRN1 and its distinctive role in chlorophyll breakdown during leaf senescence in bamboo (Bambusa emeiensis ‘Viridiflavus’). Plant Cell, Tissue and Organ Culture, 114(1), 1–10. https://doi.org/10.1007/s11240-013-0298-y

Weigel, D., Ahn, J. H., Blázquez, M. A., Borevitz, J. O., Christensen, S. K., Fankhauser, C., Ferrándiz, C., Kardailsky, I., Malancharuvil, E. J., Neff, M. M., Nguyen, J. T., Sato, S., Wang, Z. Y., Xia, Y., Dixon, R. A., Harrison, M. J., Lamb, C. J., Yanofsky, M. F., & Chory, J. (2000). Activation tagging in Arabidopsis. Plant Physiology, 122(4), 1003–1013. https://doi.org/10.1104/pp.122.4.1003

Yang, R., Howe, J. A., & Golden, B. R. (2018). Calcium silicate slag reduces drought stress in rice (Oryza sativa L .). June, 353–361. https://doi.org/10.1111/jac.12327

Yin, X., Huang, L., Zhang, X., Guo, C., Wang, H., Li, Z., & Wang, X. (2017). Expression patterns and promoter characteristics of the Vitis quinquangularis VqSTS36 gene involved in abiotic and biotic stress response. Protoplasma, 254(6), 2247–2261. https://doi.org/10.1007/s00709-017-1116-x

Yoine, M., Nishii, T., & Nakamura, K. (2006). Arabidopsis UPF1 RNA helicase for nonsense-mediated mRNA decay is involved in seed size control and is essential for growth. Plant and Cell Physiology. https://doi.org/10.1093/pcp/pcj035

Yoshida, T., Mogami, J., & Yamaguchi-Shinozaki, K. (2014). ABA-dependent and ABA-independent signaling in response to osmotic stress in plants. In Current Opinion in Plant Biology (Vol. 21, pp. 133–139). Elsevier Ltd. https://doi.org/10.1016/j.pbi.2014.07.009

Yunes, J. A., Neto, G. C., da Silva, M. J., Leite, A., Ottoboni, L. M. M., & Arruda, P. (1994). The transcriptional activator Opaque2 recognizes two different target sequences in the 22kD-like alpha-prolamin genes. Plant Cell, 6(2), 237–249. https://doi.org/10.1105/tpc.6.2.237

Zhang, C., Shi, S., Wang, B., & Zhao, J. (2018). Physiological and biochemical changes in different drought 17 tolerant alfalfa (Medicago sativa L .) varieties under PEG 17 induced drought stress. Acta Physiologiae Plantarum, 40(2), 1–15. https://doi.org/10.1007/s11738-017-2597-0

Zhang, J., Jia, W., Yang, J., & Ismail, A. M. (2006). Role of ABA in integrating plant responses to drought and salt stresses. Field Crops Research, 97(1 SPEC. ISS.), 111–119. https://doi.org/10.1016/j.fcr.2005.08.018

Zhou, S., Zhang, H., Li, R., Hong, Q., Li, Y., Xia, Q., & Zhang, W. (2017). Function identification of the nucleotides in key cis-element of Dysfunctional Tapetum1 (DYT1) promoter. Frontiers in Plant Science, 8(February), 1–8. https://doi.org/10.3389/fpls.2017.00153

Zhu, M., Chen, G., Dong, T., Wang, L., Zhang, J., & Zhao, Z. (2015). SlDEAD31, a Putative DEAD-Box RNA Helicase Gene, Regulates Salt and Drought Tolerance and Stress-Related Genes in Tomato. 1–20. https://doi.org/10.1371/journal.pone.0133849

