## Supplementary file for "Gain of function mutagenesis through activation tagging identifies *XPB2* and *SEN1* helicase genes as potential targets for drought stress tolerance in rice"

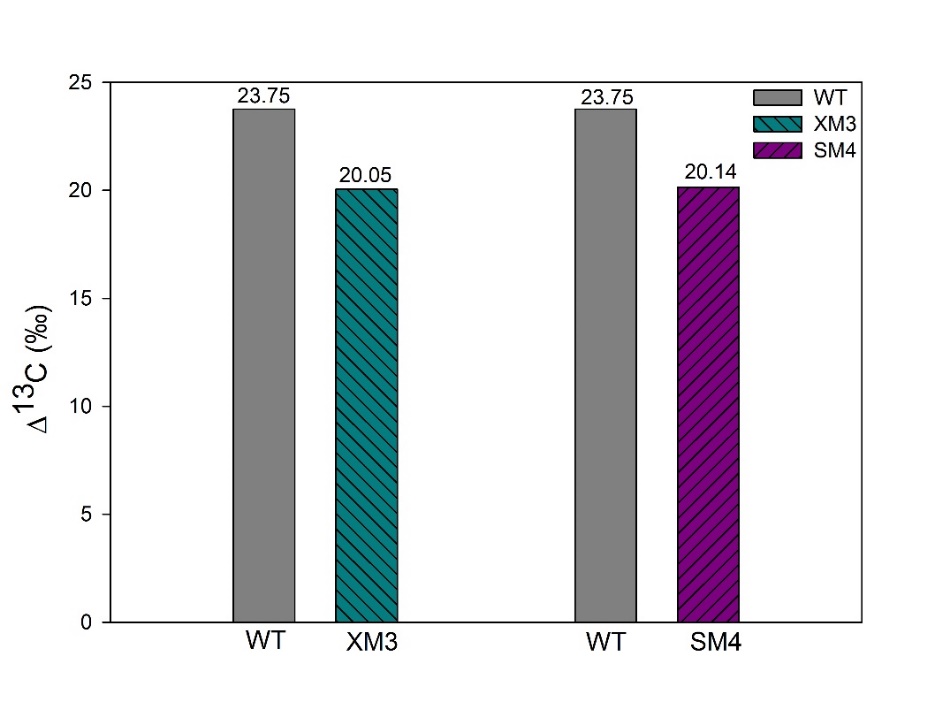

Figure S1: Carbon isotope discrimination (Δ^13^C‰) as measured by isotope ratio mass spectrometer. This is an indirect method of identifying plants with higher WUE under water stress conditions. A high Δ^13^C value indicates a lower WUE. Here the WT lines were observed to have 23.75‰ of Δ^13^C under limited water conditions which was higher than that of XM3 (20.05‰) and SM4 (20.14‰) lines.

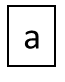

Figure S2 :(a) *In-silico* analysis of 1Kb upstream promoter region of SEN1. The sequence was retrieved from rice genome database and was subjected to PlantCARE online search tool for locating the elements. Each element has been colour coded and the index is provided along with the figure.

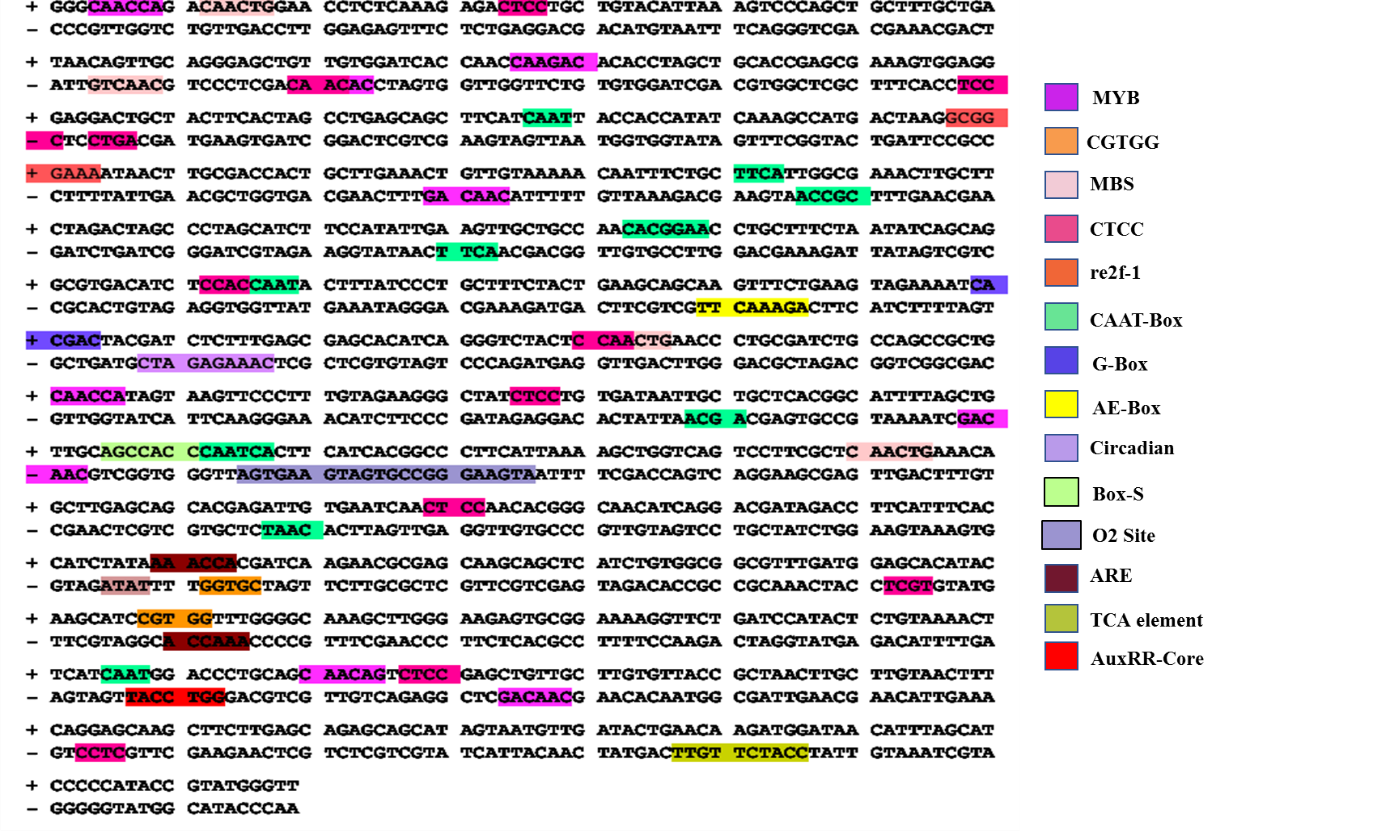

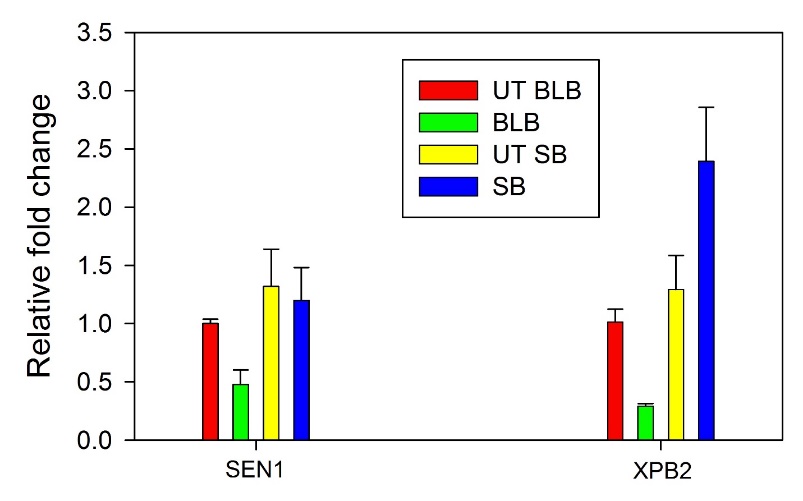

Figure S3. : Quantitative real-time PCR (qRT-PCR) analyses of *SEN1* and *XPB2* genes in response to *Xanthomonas* *oryzae* pv. *oryzae* (*Xoo*) and *Rhizoctonia* *solani,* which cause Bacterial Leaf Blight (BLB) and Sheath Blight (SB) respectively. The data was normalized using untreated (UT) plant samples grown under similar conditions. Not much upregulation was observed under biotic stress conditions. The fold change was calculated using ΔΔC_T_ method.

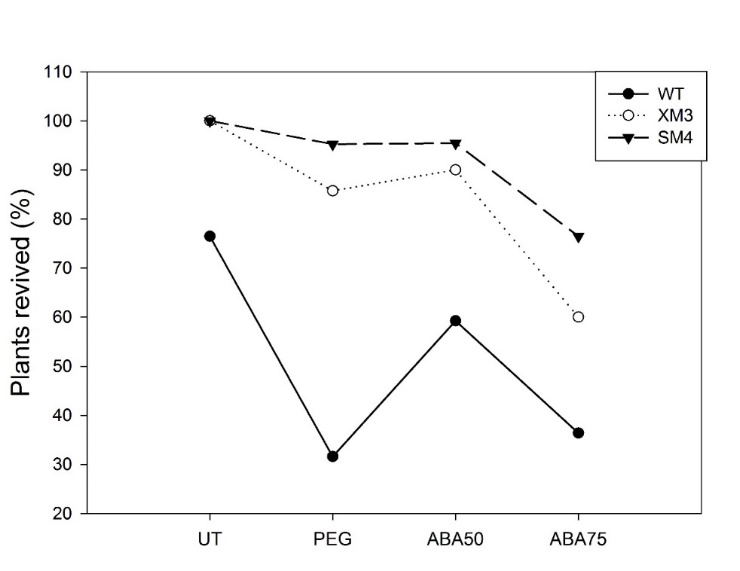

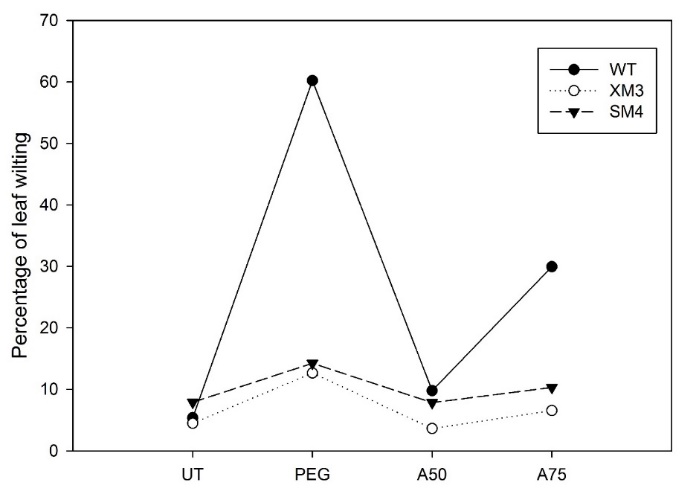

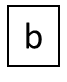

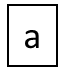

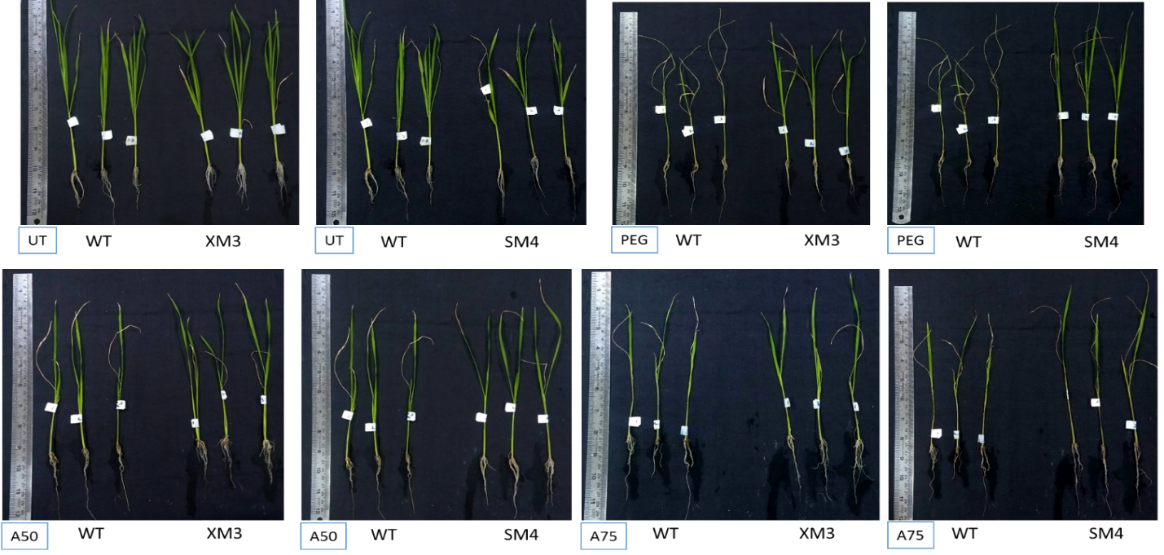

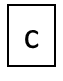

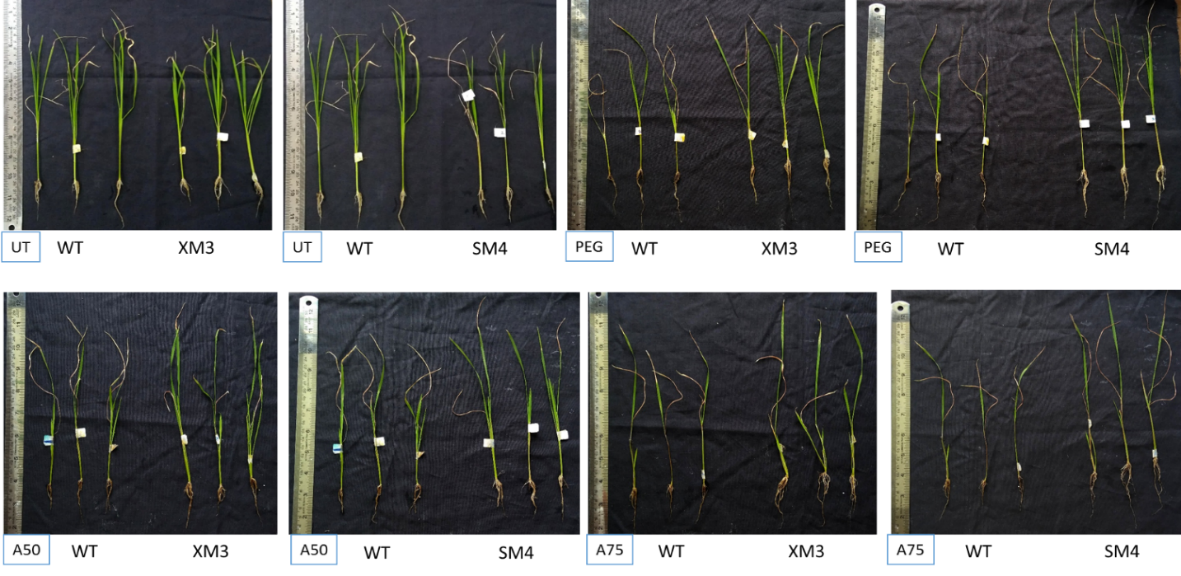

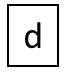

**Figure S4:** Depicting the phenotypes, wilting percentage and recovery of the plants WT,XM3 and SM4 lies 20DAS and 20 DAR. (a) shows the leaf wilting percentage 20 DAS and (b) depicting the percentage of revival of plant 20 DAR; (c) and (d) showing phenotypic difference of XM3 and SM4 20 DAS and 20 DAR respectively in comparison to wild type. The wild type plants experienced a very high rate of wilting (60%) and low rate of revival (30%) under PEG treatment as compared to the tagged lines. XM3 and SM4 showed 12%-14% wilintg under PEG and more than 85% revival was observed under both PEG and 50µM ABA. The stress conditions imposed were 10% PEG, 50µM ABA and 75µM ABA.

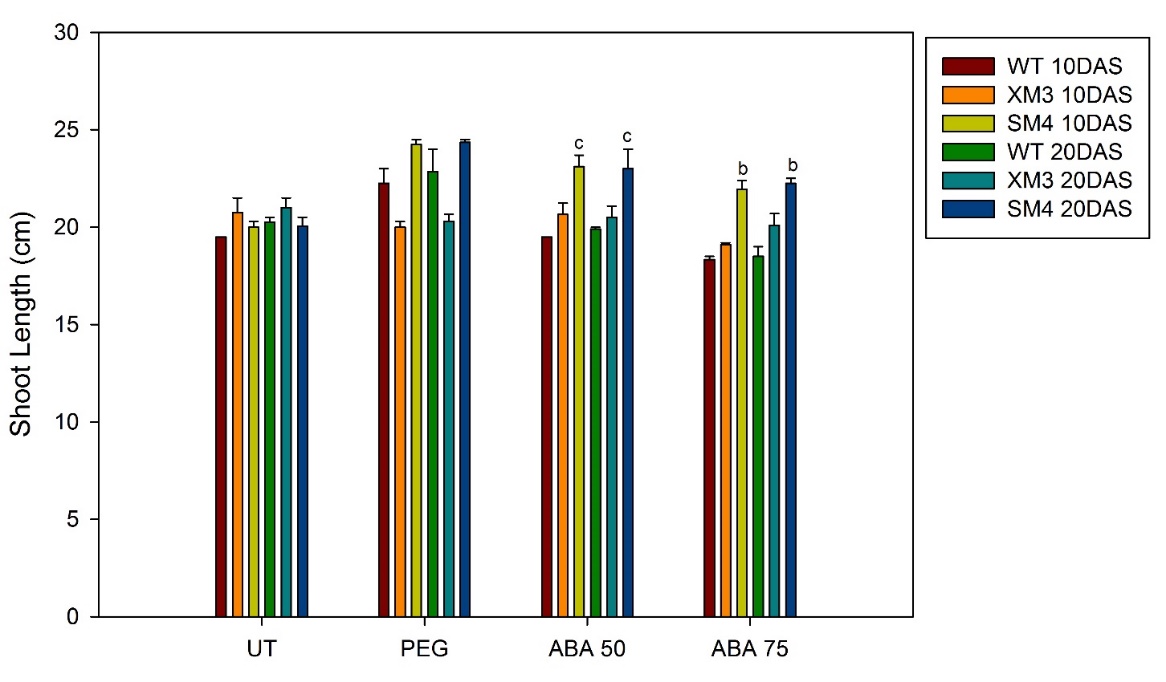

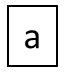

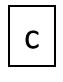

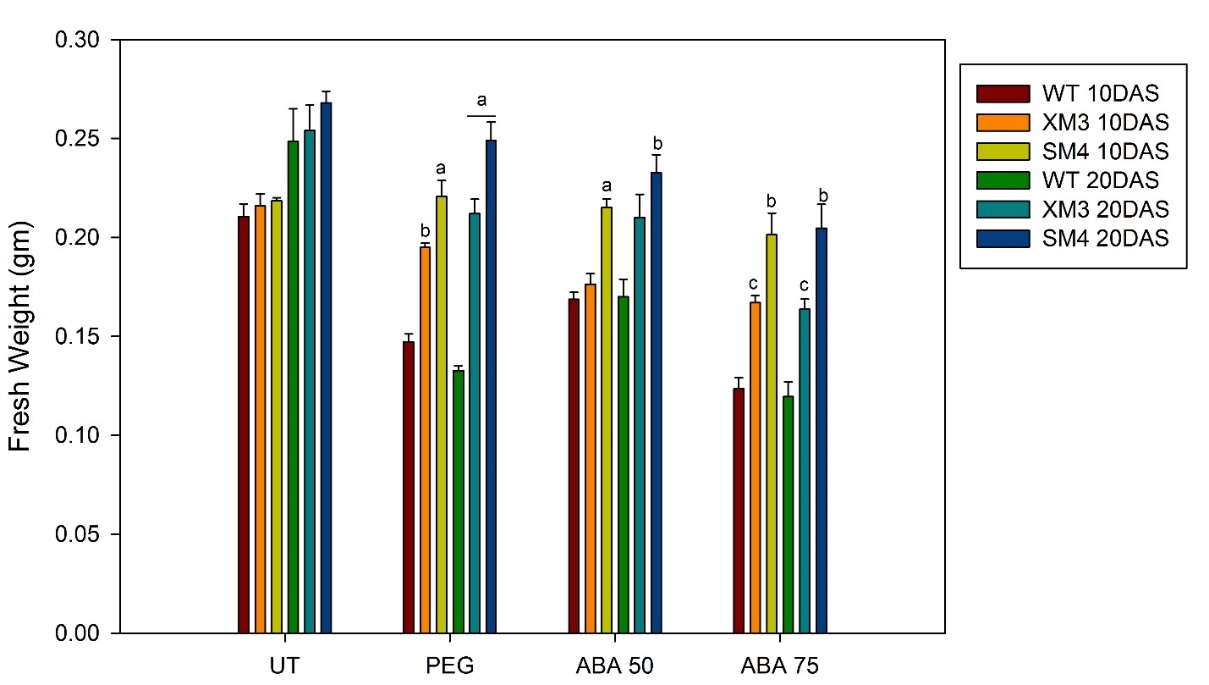

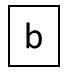

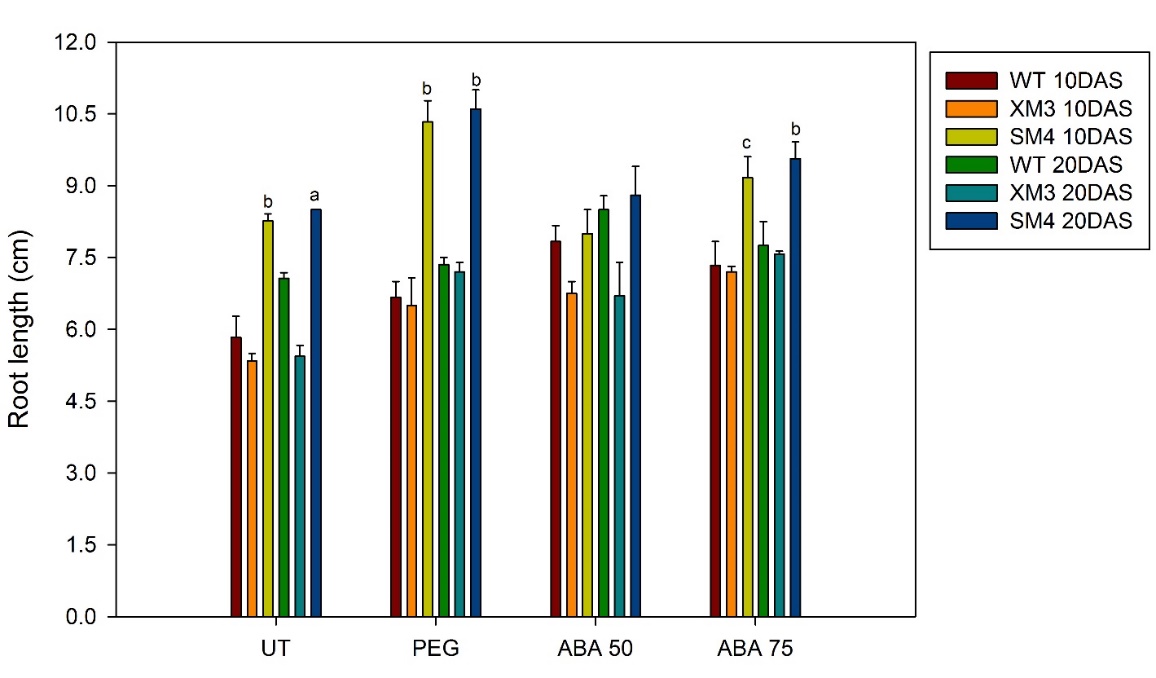

Figure: S5: Growth parameters of XM3 and SM4 in comparison to the wild type plants under various imposed stress conditions. The mean of the readings taken 10 and 20 DAS were plotted in a bar chart. (a) Shows the variation in the fresh weight; (b) and (c) shows the variation in the shoot and root length respectively. A higher fresh weight was maintained by XM3 and SM4 under stress conditions in contrary to the WT plants. The shoot length did not differ much in between the tagged lines and the WT plants, but, the root length of SM4 lines were observed to be significantly high under stress as compared to the WT plants. The mean and the standard error is plotted in a vertical bar graph. One way ANOVA was performed at a significance level P<0.001 marked as a, P<0.025 marked as b and P<0.05 marked as c.

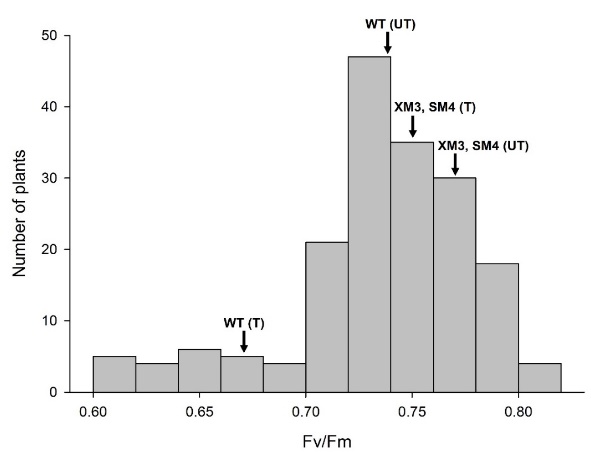

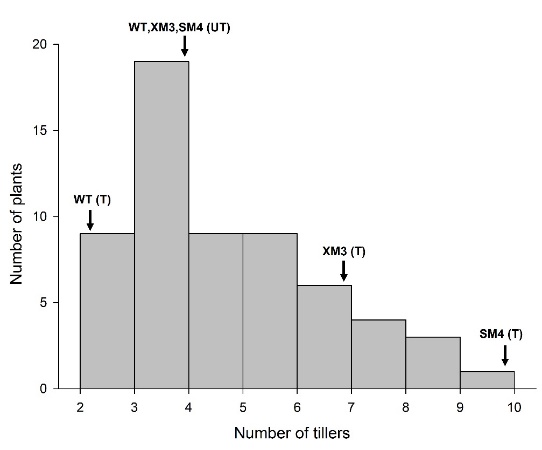

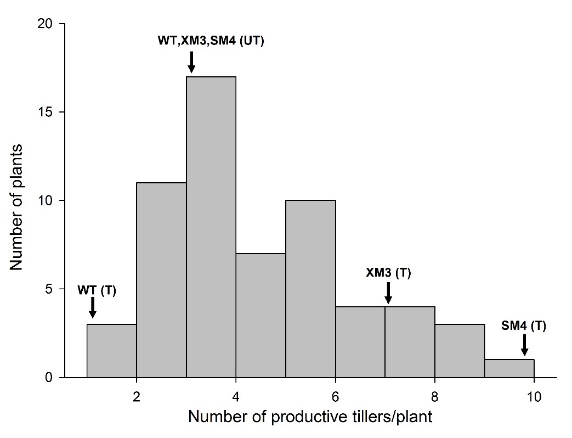

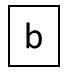

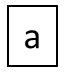

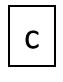

Figure
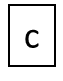
S6: Physiological analysis of the tagged lines. (a) and (b) represents number of tillers/plant and number of productive tillers/plant which ranged from 3-9/plant, all of them being productive in the tagged lines. The WT had 2-3 tillers/plant, 1-2 of them were productive upon stress imposition.(UT-untreated; T-treated). The readings are depicted in the form of a histogram. (c) Depicts the photosynthetic efficiency as measured by the mini PAM of the tagged lines in comparison with the wild type plants. The Fv/Fm ratio was 0.72 to 0.77 in XM3 and SM4 even after application of stress as compared to the wild type plants whose efficiency dropped to 0.63 to 0.67. All the values have been plotted as a histogram.

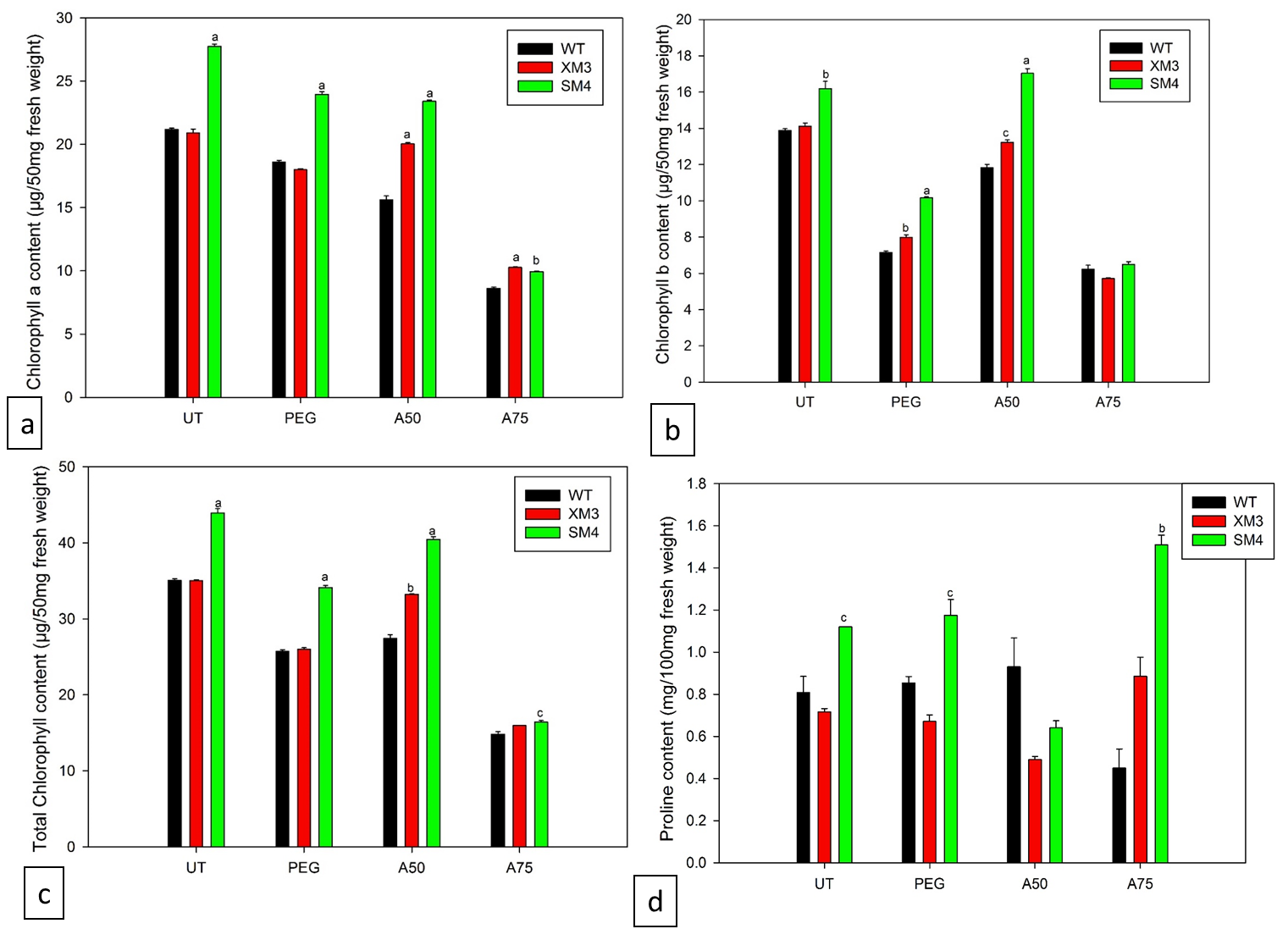

Figure S7: Graphical representation of the biochemical studies done on XM3 and SM4 in comparison to WT plants post stress. (a), (b), (c), (d) depicts chlorophyll a, chlorophyll b, total chlorophyll and proline content post revival respectively. One way ANOVA was performed at a significance level P<0.001 marked as a, P<0.025 marked as b and P<0.05 marked as c.

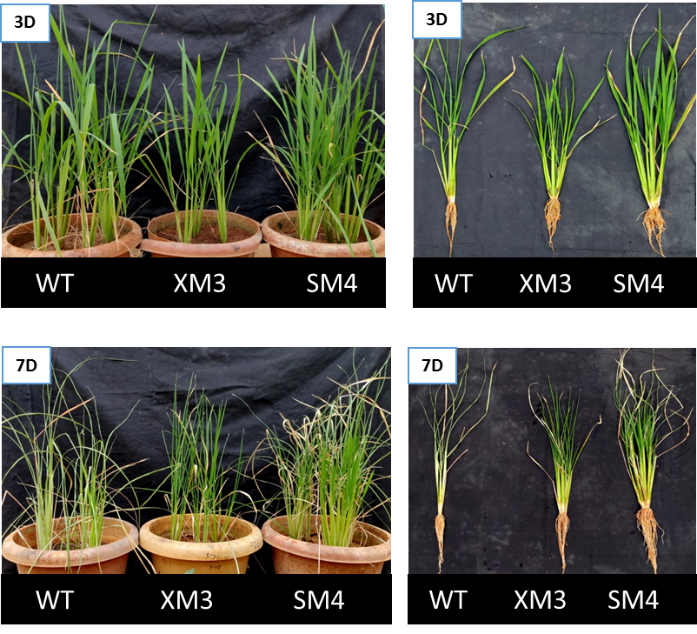

Figure S8: Phenotypic observation of the WT, XM3 and SM4 plants after periodic drought of 3 and 7 days. The WT lines showed less green phenotype with more leaf curling and wilting and poor root development.

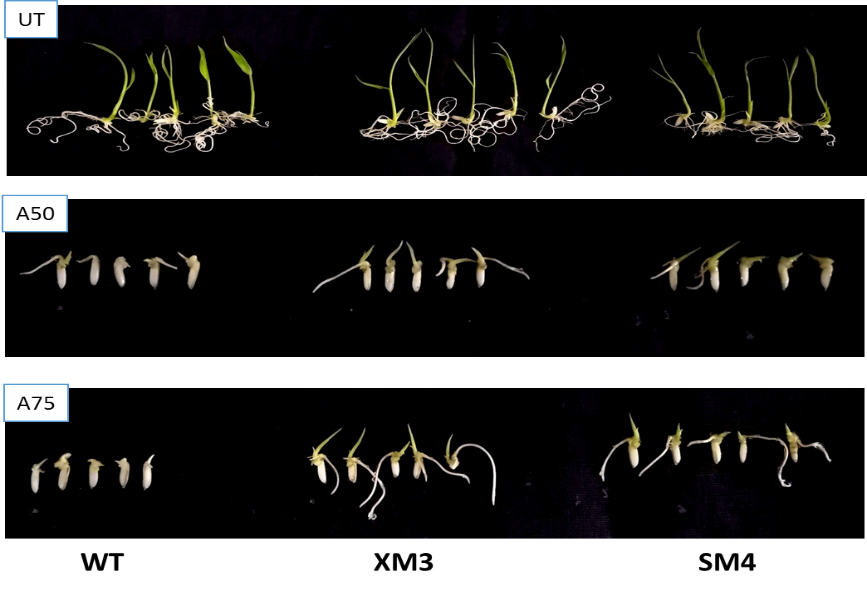

Figure S9: Depicting the seedling germination after 5 days. A growth retardation in the WT seedlings was observed under stress conditions (ABA 50µM and ABA 75µM) but the tagged lines continued to grow under high ABA concentration

**Table S1:** List of cis-acting elements and their functions found in the 1Kb upstream promoter region of SEN1 through PlantCARE database.

| Element | Position | Strand | Sequence | Function |
| --- | --- | --- | --- | --- |
| AE-Box | 399 | - | AGAAACTT | Light response module (Wei et al., 2013) |
| ARE | 709 | + | AAACCA | Element for anaerobic induction (Wei et al., 2013) |
|  | 780 | - | AAACCA |  |
| AuxRR Core | 847 | - | GGTCCAT | Element for auxin response (Sakai et al., 1996) |
| CAAT Box | 176 | + | CAAT | Element in promoter and enhancer region |
|  | 366 | + | CAAT |  |
|  | 365 | + | CCAAT |  |
|  | 647 | - | CAAT |  |
|  | 264 | - | CCAAT |  |
|  | 571 | + | CCAAT |  |
|  | 317 | + | TGCCAAC |  |
|  | 845 | + | CAAT |  |
|  | 251 | + | CAAT |  |
|  | 536 | - | CAAT |  |
|  | 306 | - | CAAT |  |
|  | 572 | + | CAAT |  |
| G-box | 419 | + | CACGAC | Light responsivenes element ( Wei et al., 2013) |
| MBS | 12 | + | CAACTG | MYB binding site for drought inducibility (Ambawat et al., 2013) |
|  | 461 | + | CAACTG |  |
|  | 74 | - | CAACTG |  |
|  | 620 | + | CAACTG |  |
| MYB | 4 | + | CAACCA |  |
|  | 491 | + | CAACCA |  |
|  | 239 | - | CAACCA |  |
|  | 860 | + | CAACCA |  |
|  | 87 | - | CAACCA |  |
|  | 558 | - | CAACCA |  |
|  | 101 | + | CAACCA |  |
|  | 874 | - | CAACCA |  |
| O2- site | 575 | - | GATGA(C/T)(A/G)TG(A/G) | Element involved in zein metabolism induction (Yunes et al., 1994) |
| TATA- box | 705 | - | TATA | Core promoter element |
| TCA-element | 957 | - | CCATCTTTTT | Element for salicylic acid responsiveness (Wei et al., 2013) |
| Unnamed_1 | 712 | - | CGTGG | ABRE motif (Srivastav et al., 2010) |
|  | 778 | + | CGTGG |  |
| Unnamed_4 | 34 | + | CTCC | Regulates gene expression for anther development (Zhou et al., 2017) |
|  | 525 | + | CTCC |  |
|  | 360 | + | CTCC |  |
|  | 913 | - | CTCC |  |
|  | 136 | - | CTCC |  |
|  | 760 | - | CTCC |  |
|  | 458 | + | CTCC |  |
|  | 659 | + | CTCC |  |
|  | 83 | - | CTCC |  |
|  | 867 | + | CTCC |  |
|  | 140 | - | CTCC |  |
| Box S | 565 | + | AGCCACC | Responsive to wounding and pathogen ellicitation (Yin et al., 2017); Stress responsiveness (Ding et al., 2019) |
| Circadian | 428 | - | CAAAGATATC | Element for circadian control (Fu et al., 2014) |
| re2f-1 | 207 | + | GCGGGAAA | E2F binding site ( Chabouté et al., 2002); Responsible for cell cycle (Çakir et al.,2013) |

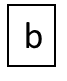

Table S2: Chart showing phenotypic characteristics observed in the tagged lines and the WT plants post acclimatization in the greenhouse. The observations included number of primary branch/panicle, numbers of seeds/branch of the panicle, seeds/panicle, total seeds/plant, boot leaf length and the panicle length. The mean ± standard error is represented in the chart. One way ANOVA was performed at a significance level P<0.001 marked as a, P<0.025 marked as b and P<0.05 marked as c.

| Treatment | Parameter | WT | XM3 | SM4 |
| --- | --- | --- | --- | --- |
| UT | Primary branch/panicle | 7.33 ± 0.66 | 7.66 ± 0.33 | 7.33 ± 0.33 |
|  | Seeds/branch | 15.55 ± 1.46 | 17.4667 ± 2.25 | 17.1538 ± 1.58 |
|  | Seeds/panicle | 104.33 ± 5.2 | 102.66 ± 7.17 | 116.33 ± 9.24 |
|  | Total seeds/plant | 328 ± 21.21 | 299.5 ± 8.5 | 324.5 ± 24.5 |
|  | 100 seeds weight (g) | 1.30 ± 0.02 | 1.28 ± 0.3 | 1.48 ± 0.01^a^ |
|  | Boot leaf length (cm) | 22.28 ± 1.04 | 21.62 ± 1.4 | 27.71 ± 1.13^b^ |
|  | Panicle length (cm) | 13.62 ± 0.66 | 13.42 ± 0.69 | 14.79 ± 0.43 |
|  | Plant height (cm) | 73.25±1.06 | 68±1.15 | 78.4±0.97 |
| PEG 10% | Primary branch/panicle | 4.28±0.28 | 5.7±0.22^b^ | 6.5±0.22^a^ |
|  | Seeds/branch | 8.62±.420 | 12.52±.442^a^ | 14.16±.3^a^ |
|  | Seeds/panicle | 35.2±1.42 | 69.25±5.15^a^ | 84.12±4.21^a^ |
|  | Total seeds/plant | 68.5 ± 0.5 | 277 ± 30^b^ | 349.5 ± 31.5^b^ |
|  | 100 seeds weight (g) | 1.24 ± 0.02 | 1.36 ± 0.00^a^ | 1.45 ± 0.01^a^ |
|  | Boot leaf length (cm) | 18.7 ± 0.86 | 23.9 ± 0.8^b^ | 28.11 ± 0.65^a^ |
|  | Panicle length (cm) | 10.92 ± 0.54 | 14.43 ± 0.5^a^ | 15.99 ± 0.35^a^ |
|  | Plant height (cm) | 49.5±1.5 | 57.1±1.64 | 64.5±1.50^a^ |
| ABA 50µM | Primary branch/panicle | 5.6±0.4 | 4.3±.21^c^ | 7.1±.26^b^ |
|  | Seeds/branch | 10.3±.365 | 10.3±.579 | 16.64±.494^a^ |
|  | Seeds/panicle | 50.44±.365 | 45±4.88 | 111.62±4.72^a^ |
|  | Total seeds/plant | 151.33 ± 12.7 | 206.5 ± 18.5 | 446.5 ± 7.5^a^ |
|  | 100 seeds weight (g) | 1.20 ± 0.04 | 1.39 ± 0.016^b^ | 1.42 ± 0.03^b^ |
|  | Boot leaf length (cm) | 18.83 ± 0.83 | 20.32 ± 0.61 | 30.33 ± 0.75^a^ |
|  | Panicle length (cm) | 13.08 ± 0.57 | 14.36 ± 0.37 | 17.02 ± 0.35^a^ |
|  | Plant height (cm) | 47±1 | 52±1.15 | 72.25±1.18^a^ |
| ABA 75µM | Primary branch/panicle | 1±0 | 5.33±.210^b^ | 7.16±.477^a^ |
|  | Seeds/branch | 11±0 | 14.17±.565 | 13.83±.382 |
|  | Seeds/panicle | 11±0 | 69±7.17^b^ | 98.93±6.83^b^ |
|  | Total seeds/plant | 11±0 | 461 ± 16^b^ | 557.5 ± 5.5^b^ |
|  | 100 seeds weight (g) | 0.116 ± 0 | 1.35 ± 0.04^a^ | 1.34 ± 0.03^a^ |
|  | Boot leaf length (cm) | 11 ± 0 | 25.21 ± 1.34^b^ | 29.8 ± 1.25^b^ |
|  | Panicle length (cm) | 9 ± 0 | 16.29 ± 0.42^a^ | 17.13 ± 0.38^a^ |
|  | Plant height (cm) | 28±0 | 56±3^c^ | 78.5±0.5^b^ |

Table S3: List of primers used

| Name of Primer | 5’ to 3’ sequence | Purpose used for |
| --- | --- | --- |
| SEN1 RT | AATCATGGTGTGGGTTTCGT | Differential transcript analysis |
| XPB2 RT | TCAATGGGCATTTCAGTTCA | Differential transcript analysis |
| DS.SP1 | CTCACAGCACTTAGCAGTACAGCACGTCAGC | Nested primer in Primary TAIL PCR |
| DS.SP2 | GTGCGCGTGGGCATGGATGTGGC | Nested primer in Secondary TAIL PCR |
| DS.SP3 | ATAGTTTAGTTAAAGGTCAGTTGTGTC | Nested primer in Tertiary TAIL PCR |
